# High quality *de novo* genome assembly of non-conventional yeast *Kazachstania bulderi* a new potential low pH production host for biorefineries

**DOI:** 10.1101/2023.01.11.523663

**Authors:** L. N. Balarezo-Cisneros, S. Timouma, A. Hanak, A. Currin, F. Valle, D Delneri

**Affiliations:** Manchester Institute of Biotechnology, University of Manchester, Manchester, UK; BP Biosciences Center, San Diego, California, USA

## Abstract

*Kazachstania bulderi* is a yeast species belonging to a ubiquitous group of non-conventional yeasts which has the ability to grow efficiently on glucose and δ-gluconolactone at low pH. This unique trait makes *K. bulderi* an ideal candidate as a new host for low pH fermentation processes for sustainable production of green chemicals such as organic acids. To accelerate strain development with this species, detailed information of its genetics is needed. Here, by employing high accuracy long read sequencing we report a high-quality phased genome assembly for three strains belonging to *K. bulderi* species, including the type strain. The sequences were assembled into 12 chromosomes with a total length of 14Mb, and the genome was fully annotated at structural and functional levels, including allelic and structural variants, ribosomal array, centromeres and mating type locus. This high-quality reference genome provides an essential resource to advance our fundamental knowledge of biotechnologically relevant non-conventional yeasts and to support the development of genetic tools for manipulating such strains towards their use as production hosts biotechnological processes.

## Introduction

Biorefineries have been proposed as a solution to replace oil-derived products with more sustainable biotechnologies, which are based on the use of renewable resources for the commercial production of chemicals and other products. These types of processes are often hampered by the high costs associated with the recovery and purification of the product(s), which in many cases, can be as high as 50%-80% of the total production cost (1–3) This is particularly evident for the production of organic acids by fermentation. Because most production microorganisms used in the industry need a neutral pH for optimal performance during the fermentation process, as the organic acids are synthesized and accumulated in the culture media, pH decreases, and the organic acid has to be neutralized by the addition of a base, leading to the formation of the organic acid salt (4,5). Subsequent applications of the organic acids normally use the acid form, and the neutralizing cation has to be removed, disposed or recycled (6). There is, therefore, a need to develop new production hosts that have an optimum performance at low pH. Production of organic acids at low pH, would decrease or eliminate the formation of the organic salts, and improve product recovery and overall process economics. Three strains of *Kazachstania bulderi*, formerly *Saccharomyces bulderi*, isolated from corn silage (7), have been reported to rapidly and efficiently, ferment glucose and δ-gluconolactone to ethanol and carbon dioxide at pH between 2.5 and 5.0 (8), making *K. bulderi* a very attractive host for low-pH fermentations. However, the development of a new production host for commercial processes, requires a deep understanding of its genomic sequence and organization, with accurate gene annotations, as well as the development of genetic tools to modify the strain to create the desired phenotypes.

*Kazachstania*, as their closer neighbours, *Naumovozyma* and *Saccharomyces* genera, arose after the whole-genome duplication event that gave rise to part of the *Saccharomycetaceae* (9). *Kazachstania* genus encompasses a large and diverse group of ascomycetous budding yeasts. To date, it is composed of over 40 species isolated from wild, domesticated and clinical environments (7,10).

Currently, high quality genome assemblies at chromosome level for *Kazachstania* strains are still limited, as only four assembled and annotated genomes have been published to date for *K. africana* (11), *K. naganishii* (11), *K. saulgeensis* (12) and *K. barnettii* (10). Highly fragmented assemblies have been drafted for seven species, such as *K. humilis* (13), *K. servazzii* (14,15), *K. telluris* (16), *K. slooffiae* (17), *K. bovina* (18), *K. exigua* and *K. unispora* (17); other species have a draft assembly with very few, or no annotations (19,20). A fragmented genome for *K. bulderi* strain appears has been deposited in NCBI, but there is no publication attached to it.

Here, using a combination of next generation sequencing, *de novo* assembly tools and gene annotation algorithms, we construct high-quality reference genomes, including ribosomal repeats, mating type locus, and mitochondria DNA for the three reported *K. bulderi* strains CBS 8638 (type strain), CBS 8639 and NRRL Y-27205. We also carried out comparative genomic analysis of the strains, including allelic and structural variation, synteny, and identification of species-specific and genus-specific genes. The availability of this genome will not only allow the optimization of gene manipulation tools towards the development of sustainable green products but will also provide a platform for future population genomic, or ‘omics studies for *Kazachstania species*.

## Results

### 1. Phenotype characteristic of *K. bulderi* strains at low pH, lactic acid and drugs

To confirm the potential of *K. bulderi* as a new production host for low-pH fermentations, we assessed the growth of three *K. bulderi* strains under different range of pH, from 5.5 to 2.1 on minimal SD media. We observed that, in contrast to *Saccharomyces cerevisiae* lab strain BY4743 and commercial strain NCYC 505, the three *K. bulderi* strains CBS 8638, CBS 8639 and NRRL Y-27205 maintain a constant high growth rate, even at pH to 2.1 (Figure 1A). CBS 8638, in particular, seems to grow better at lower pH compared to the other strains. For *Saccharomyces cerevisiae* strains, the growth is significantly affected at pH ≤ 2.5, being completely hindered at pH 2.1 for BY4743 (Figure 1A). Since an efficient host for low pH fermentation should be able to stand high concentrations of organic acids, we assessed the growth of *K. bulderi* strains in the presence of different concentration of lactic acids up to 100 g/L. At 85 g/L lactic acid, CBS 8638 strains was still able to maintain optimal growth, while the other *Kazachstania* strains showed growth delay and the *S. cerevisiae* BY4741 could not grow (Figure 1B). Only when lactic concentration was lowered to 50g/L, we could detect some growth for *S. cerevisiae* (Figure S1). We also tested the ability of the yeast strains to grow in presence of 25mM formic acid. Only the *Kazachstania* strains were able to grow in this condition (Figure 1C). All the growth parameter for the strains tested in the different conditions are reported in Table S1.

**Figure 1.**
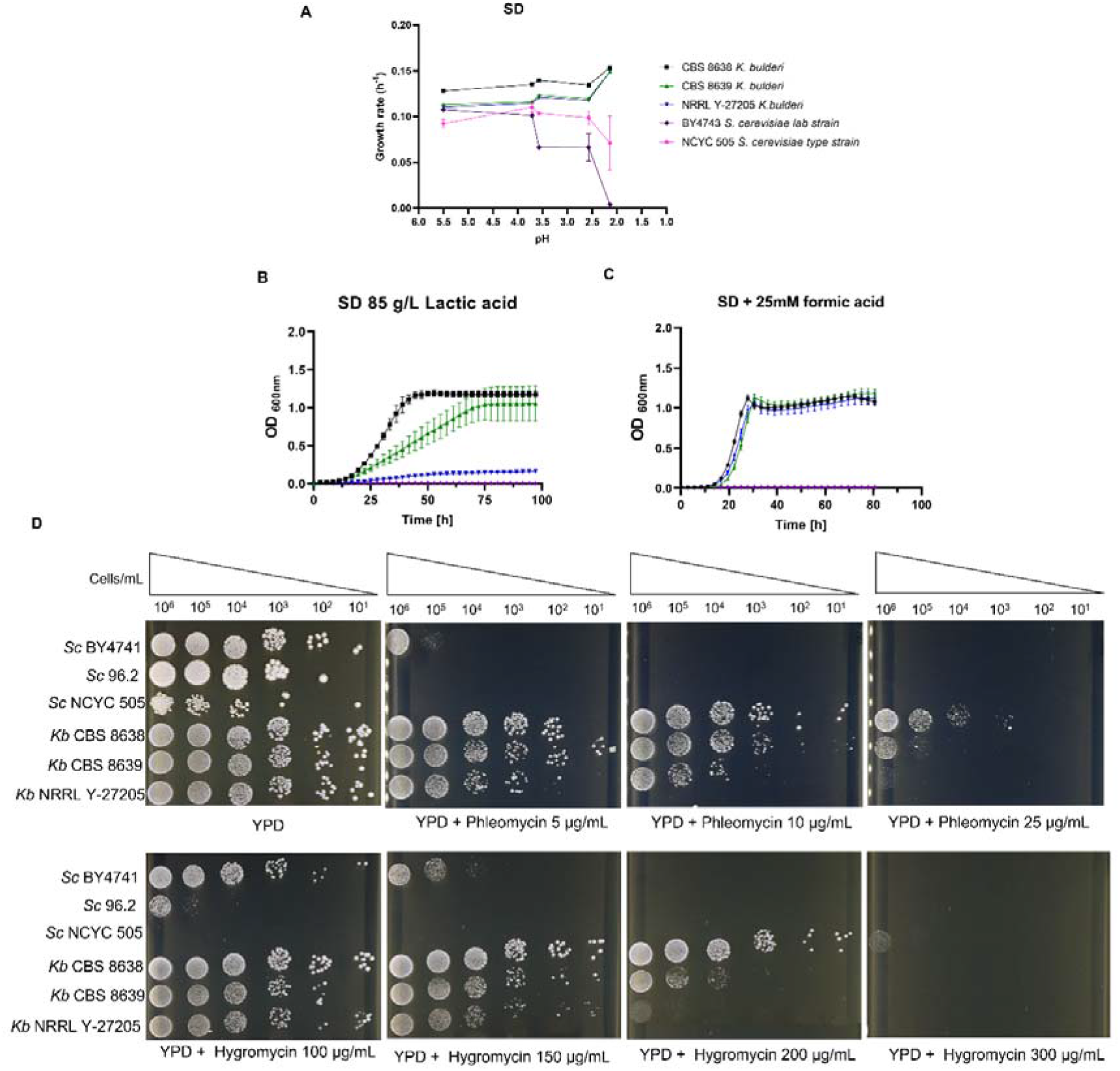
Phenotypic analysis of CBS 8638, CBS 8639 and NRRL Y-27205 *K. bulderi* strains. Panel A: maximum growth rate of the three *Kazachstania bulderi* strains compared with *Saccharomyces cerevisiae* lab BY4743 and type strain NCYC 505 from pH 5.5 to 2.5. Panel B: growth curves of *K. bulderi* strains in 85g/L of lactic acid. Panel C: growth curves of *K. bulderi* strains in 25mM of formic acid; Panel D: spot test assay of *K. bulderi* (Kb), and *S. cerevisiae* (Sc) BY4741, 96.2 and NCYC 505 strains in phleomycin and hygromycin B.

With the purpose of developing genetic tools for genetic manipulations of *K. bulderi*, we evaluated the resistance of these strains to common drugs used in the laboratory. We observed that CBS 8638, CBS 8639 showed a high resistance to the standard working concentrations of hygromycin B and phleomycin (Figure 1D) when compared with *S. cerevisiae* strains 96.2, BY4741 and NCYC 505. Interestingly, NRRL Y-27205 does not show the same resistance, being more sensible to the tested drugs. The drug that showed to be most effective for all *K. bulderi* strains was nourseothricin. These results suggest that the three tested *K. bulderi* strains display a reduce cellular uptake of cationic drugs.

### 2. Genome sequencing and *de novo* assembly of the three *K. bulderi* strains

#### a. Genome sequencing, assembly and annotation

Genome sequencing of *K. bulderi* CBS 8638, CBS 8639 and NRRL-Y27205 strains was performed using HiFi read data derived from single-molecule real-time (SMRT) technology from Pacific Biosciences (Pacbio), the *de novo* phased assembly and annotation strategies followed is summarised in Figure. S2.

We obtained approximately 131,888, 125,890 and 159,349 reads for CBS 8638, CBS 8639 and NRRL Y-27205 respectively. To assemble a high-quality genome, first we tested a combination of different phased assembly algorithms, such as the Improved Phased Assembler (IPA; the official PacBio software for HiFi genome assembly) and HIFIASM (a fast haplo-type-resolved *de novo* assembler for PacBio HiFi reads) on *K. bulderi* CBS 8638, and CBS 8639 strains. The IPA assembler consistently gave a better assembly with a number of contigs closer to the number of chromosomes found in other *Kazachstania* species (10–12), while Hifiasm assembler generated a much higher number of contigs for CBS 8638 and CBS 8639 (Table S2).

IPA generated primary *de novo* assemblies of 14, 17 and 15 contigs for CBS 8638, CBS 8639 and NRRL-Y27205 respectively, totalling *ca*. 14 Mb in length (Table S3); and alternative haplotig assemblies of 85, 108 and 172 contigs for CBS 8638, CBS 8639 and NRRL Y-27205, respectively (Table S1). These alternative haplotigs represent regions of heterozygosity, which allow separation of haplotypes for *K. bulderi* diploid strains, and spanned a total of 13Mb, 14Mb and 16 Mb (100% was separated into haplotypes). The total coverage was 59X for CBS 8638, 63Xfor CBS 8639 and 83.9X for NRRL Y-27205 (Table S3).

A high-quality assembly is determined when it respects three crucial attributes, referred as the three “C” (21): Continuous (size of contigs), Correct (how accurate) and Complete (ability to complete the whole structure of the genome). The assemblies obtained using the IPA assembler were hence selected for their better fulfilment of the three “C” criteria.

We then carried out the annotation using both AUGUSTUS (22) and YGAP (23), and the predicted proteins were functionally annotated using HybridMine (Figure 2). AUGUSTUS uses a hidden Markov model (probabilistic approach) to predict structural elements, whereas YGAP uses synteny ancestry (homology approach). HybridMine predicts one-to one ortholog (24)to infer function, and predicts groups of homologs, including paralogs, using a well annotated reference species, such as *S. cerevisiae*.

**Figure 2.**
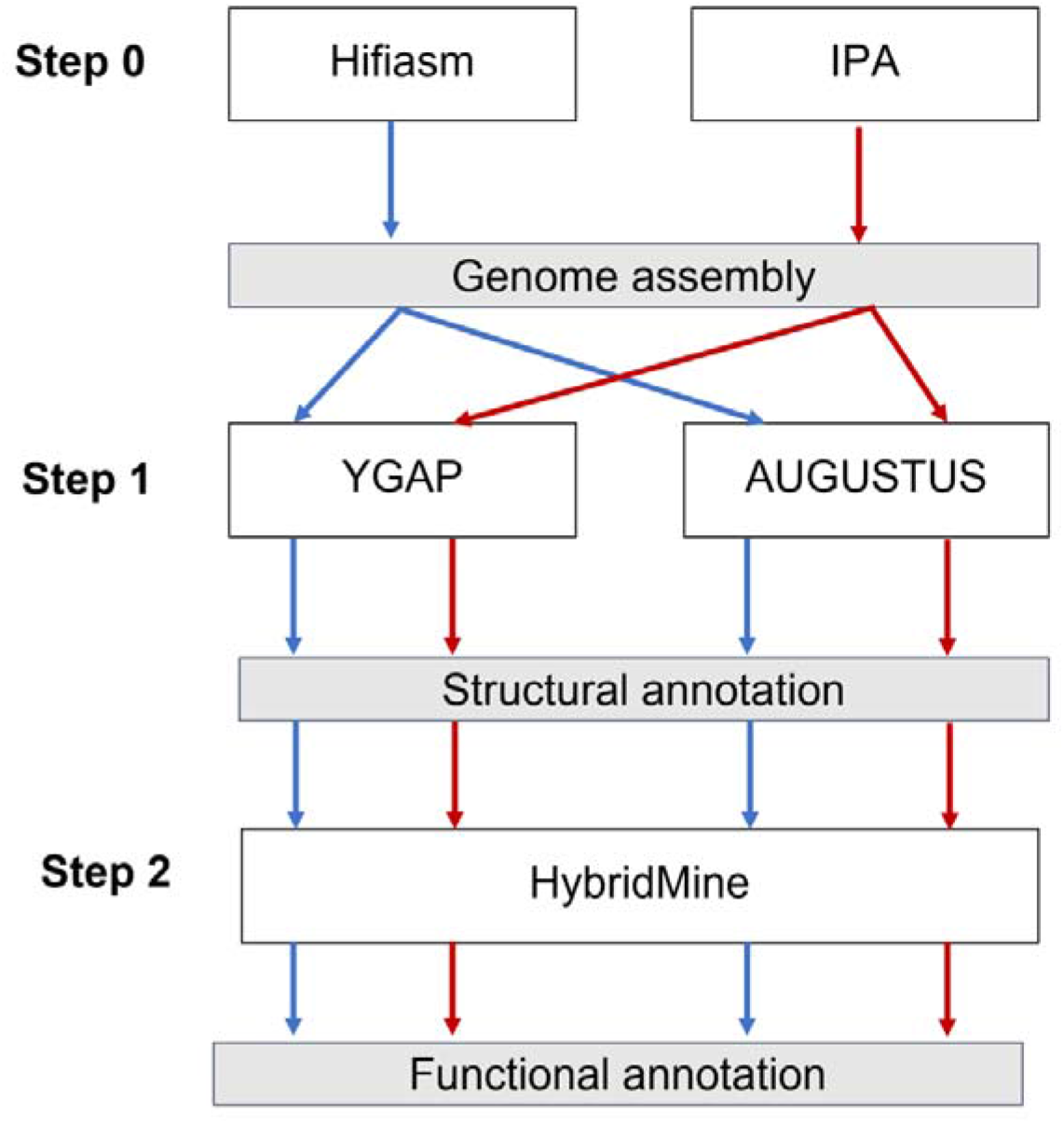
Strategy followed for generation of high-quality genome assembly of *K. bulderi* strains, including assemblers (IPA and HIFIASM), structural annotation algorithms (YGAP and AUGUSTUS) and functional annotation tool (HybridMine).

We observed that the structural annotation method that consistently produced consistently a higher number of genes functionally annotated was YGAP. AUGUSTUS predicted a higher number of genes and proteins compared to YGAP (Figure S2 A), however, when analysed with HybridMine, it also had a higher number non functionally annotated proteins, suggesting that they are either not real ORFs, or the protein sequence was not well predicted, or not translated. It is not surprising that YGAP perform better here given it is optimised towards yeast genome structural annotation. It efficiently infers introns/exons and therefore protein sequences are more accurately predicted (23)

Reliably, HybridMine was able to functionally annotate more than 90% of protein-coding genes regardless the method used (Figure S2 B).

The assembly generated using the IPA assembler and the structural annotation predicted by YGAP were used as starting point for manual curation.

#### b. Manual curation of the assemblies

When comparing individual strains assemblies between each other it was clear that: ***i***. some primary contigs in one *K. bulderi* strain were separated in two/three contigs in others; ***ii***. across the strains, there were regions with non-uniform distribution in the mapping of long reads versus the assemblies; ***iii***. translocated and inverted sequences were detected in the middle of some contigs.

The manual curation was carried out by analysing the alignment of the three assemblies of *K. bulderi*, and the mapping of HiFi reads against each assembly. Briefly, the contigs that were initially split in the different strains were verified by PCR and consolidated (*i*.*e*. now total of 12 contigs for all strains); the regions with no read coverage were troubleshooted using the alternative contig; the translocation was an artifact caused by a mis-assembly, while the inversion between CBS 8639 and NRRL Y-27205 was experimentally validated via PCR. For a detailed explanation of the curation, see the Supplementary File 1.

Following manual curation, reads were re-mapped to their respective assemblies resulting in a regular and uniform reads coverage. The primary assemblies now consist of 12 contigs, totalling 14Mb, with contig N50 of 1.2 Mb (Table 1). The coverage of the manually curated assemblies increased being now 62X, 65X and 84X for CBS 8638, CBS 8639 and NRRL Y-27205, respectively (Table 1) and a chromosome-level assembly was achieved (Table 2). *K. bulderi* CBS 8638, CBS 8639 and NRRL Y-27205 chromosomes number and their sizes were also confirmed via Pulse Field Gel Electrophoresis (Figure S3).

**Table 1.** Metrics and summary statistics for the curate *K. bulderi de novo* PacBio assemblies.

| Metric | *De novo* assemblies |  |  |
| --- | --- | --- | --- |
|  | CBS 8638 | CBS 8639 | NRRL-Y27205 |
| Size (Mb) | 14.47 | 14.25 | 14.44 |
| N° Contigs | 12 | 12 | 12 |
| Largest contig (bp) | 2767906 | 2786749 | 2758527 |
| N50 (Mb) | 1.25 | 1.24 | 1.23 |
| N75 (Mb) | 0.84 | 0.86 | 0.90 |
| L50 | 4 | 4 | 4 |
| L75 | 7 | 7 | 7 |
| GC (%) | 33.12 | 33.11 | 33.16 |
| Average identity (%) | 97.5 | 97.3 | 97.3 |
| Mean read length | 6042.8 | 6607.6 | 7153.3 |
| Median identity (%) | 99.4 | 99.3 | 99.3 |
| Median read length | 5871 | 6744 | 7308 |
| Number of reads | 149370 | 136974 | 167910 |
| Read length N50 | 7933 | 8148 | 8920 |
| Total bases | 902618975 | 905075306 | 1201106262 |
| Total bases aligned | 843227356 | 879022084 | 1162715498 |
| Coverage (X) | 62.4 | 64.5 | 83.9 |

**Table 2.**
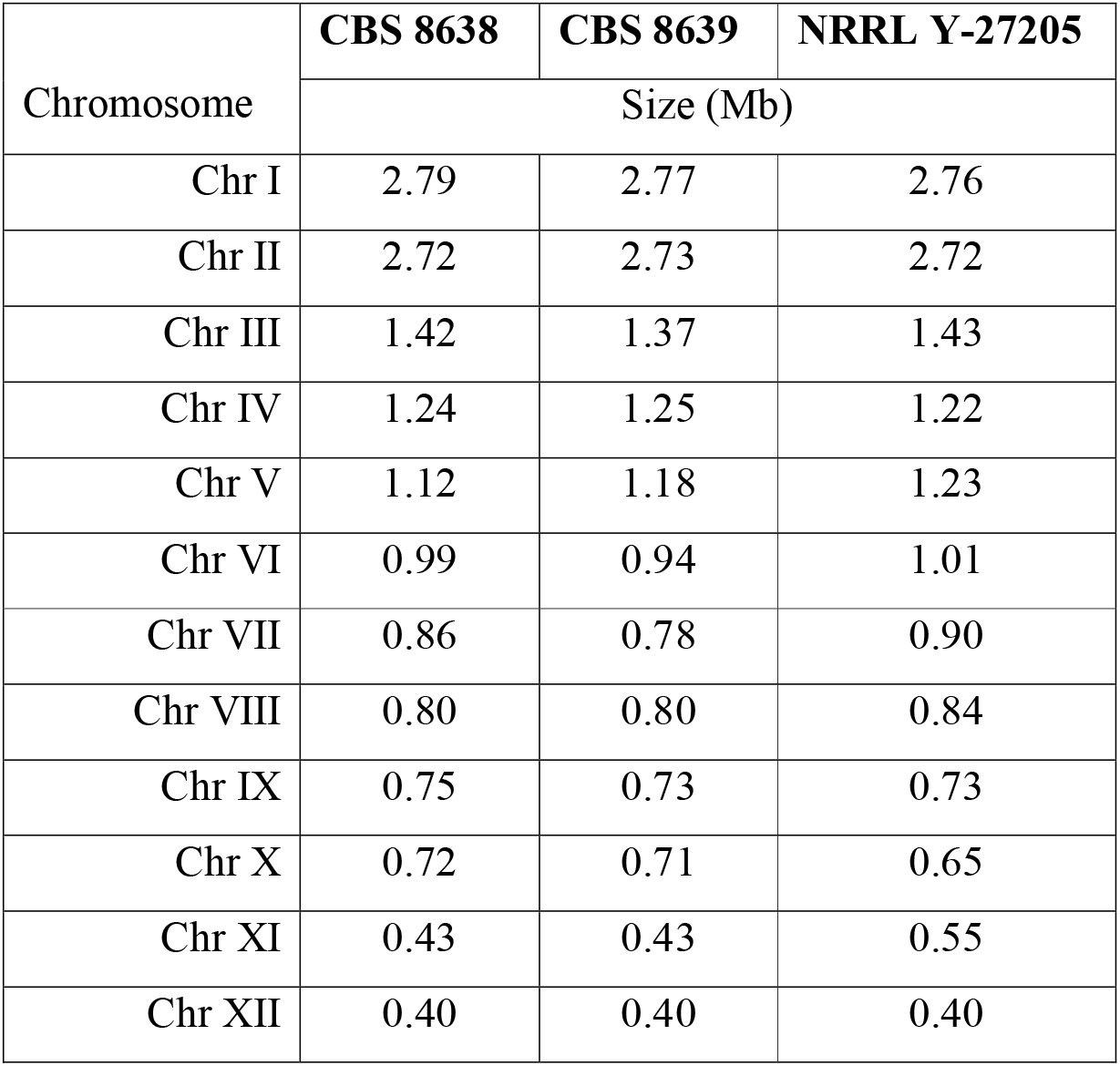
Total length of chromosome-level genome assembly for three *K. bulderi* strains after curation

The completeness of the assemblies was also evaluated using the BUSCO software using the ‘-*Saccharomycetes*’data set of BUSCO gene collection (25). Results indicate that 99% of the CBS 8638 and CBS 8639 assemblies and 98% of NRRL-Y27205 are complete (Table S4). By comparison, the previous CBS 8639 and NRRL Y-27205 assembly had 97.6% and 97.3% complete BUSCO alignments respectively. The Missing BUSCO score were 2.1% and 2.4% for both assemblies respectively, indicating a higher fraction of the genome was missing in both initial assemblies before curation (Table S4).

### 3. *K. bulderi* functional annotation

YGAP identified a total of 5877, 5759 and 5769 structural elements for CBS 8638, CBS 8639 and NRRL Y-27205, respectively, including protein coding genes and tRNAs. Furthermore, T⍰ retrotransposons and rRNAs were also annotated (Table 3, Figure 3). The genomic features annotated per chromosome, in each strain is listed in Table S5. The output of YGAP for the *K. bulderi* predicted genes used the following nomenclature: KB for *Kazachstania bulderi*; 38, 39 and Y27 for the strains CBS 8638, CBS 8639 and NRRL Y-27205, respectively; consecutive alphabet letters for the chromosome number.

**Table 3.** Annotation of the three *K. bulderi* strains.

|  | CBS 8638 | CBS 8639 | NRRL Y-27205 |
| --- | --- | --- | --- |
| Proteins coding genes | 5666 | 5551 | 5562 |
| tRNA | 211 | 208 | 207 |
| Ty | 36 | 35 | 37 |

**Figure 3.**
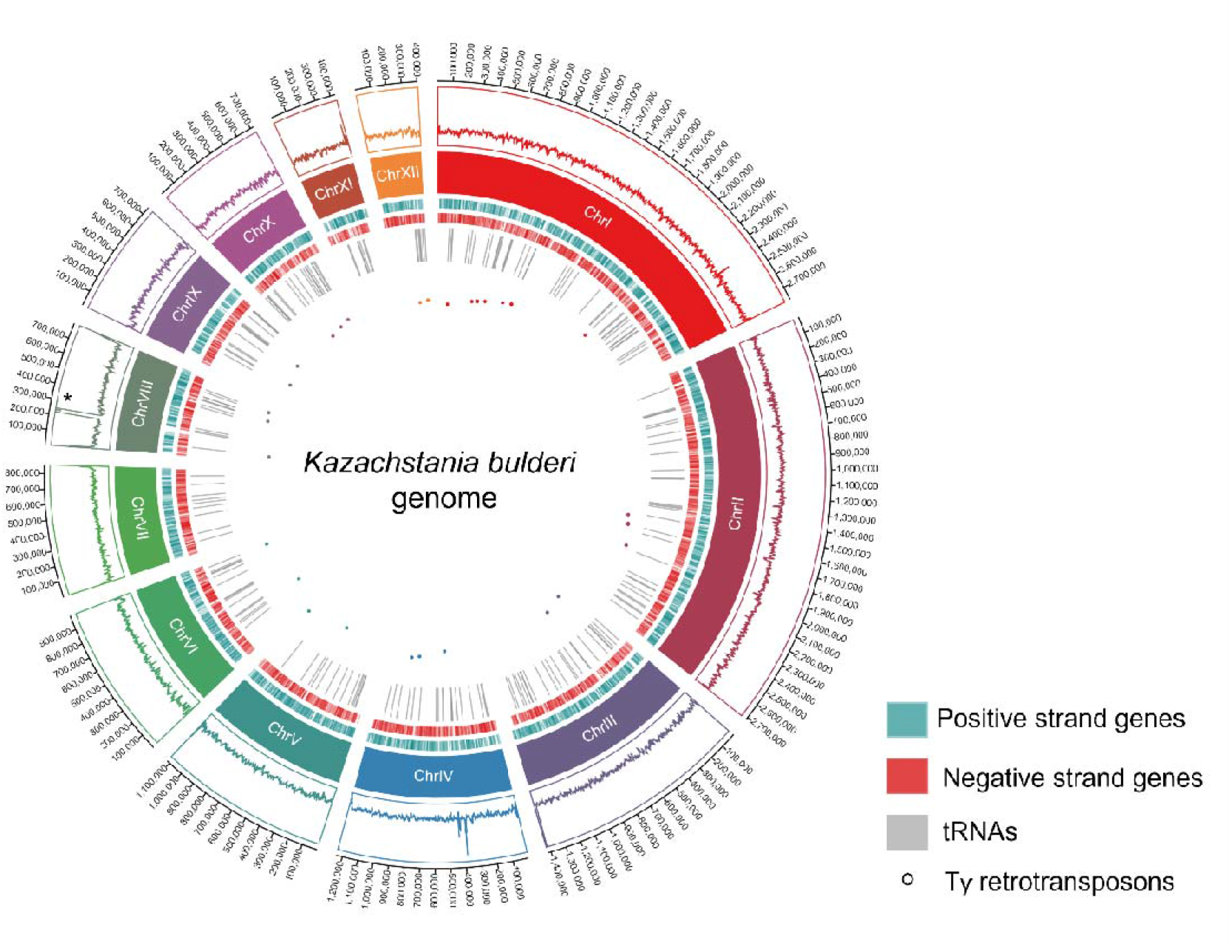
Circos Plot of 12 chromosome-level scaffolds, representing the annotation result of genes, tRNAs and transposable elements. The tracks from inside to outside are: Ty retrotransposons (round dots), tRNAs (grey), genes on positive strand (green), gene on negative strand (red), chromosomes (represented in different colours), average read coverage (reads depth track), and chromosome length. The asterisk on chromosome VIII shows the peak of reads representing the rRNA repetitions.

Functional annotation was inferred using the one-to-one orthologs found in *S. cerevisiae* model yeast. As result, 4541, 4543 and 4523 proteins had a function inferred in *K. bulderi* CBS 8638, CBS 8639 and NRRL Y-27205 respectively. Additionally, groups of homolog proteins were also identified in each strain (Dataset S1).

We searched for genes specific to the *K. bulderi*, which could not be detected by using *S. cerevisiae* as model organism. As *K. bulderi* has been isolated in maize silage, a low pH environment, it is indeed possible that these strains acquired a number of novel genes as adaptation response. To identify them, we first searched the one-to-one orthologs between *K. bulderi* and other model yeast species such as *Schizosaccharomyces pombe, Candida albicans, Candida glabrata* and *Yarrowia lipolytica* which are all well functionally annotated. Secondly, we searched one-to-one orthologs in other species that grow well at low pH, such as *Kluyveromyces marxianus, Kluyveromyces lactis, Kazachstania exigua* and *Kazachstania barnettii*, for which only structural annotations are available. In this way, we could expand the pool of annotated proteins, potentially including pH related genes, and detected genus-specific ones.

As result, a total of 5385, 5306 and 5281 proteins for CBS 8638 (Figure S4A, Dataset S2), CBS 8639 (Figure 4) and NRRL Y-27205 (Figure S4B), respectively, accounting for ~95% of all predicted proteins, had a one-to-one ortholog found in at least one of the model species used, thus they are not *K. bulderi* specific (Figure 4, Table S6). As expected, the highest number of one-to-one orthologs was detected when comparing the predicted proteins with species from *Kazachstania genus* (Table S6, Dataset S2). The 286 one-to-one orthologs between *K. bulderi* CBS 8639, *K. exigua and K. barnettii* that are not shared with the other yeast species may point out to genus specific elements (Dataset S2).

**Figure 4.**
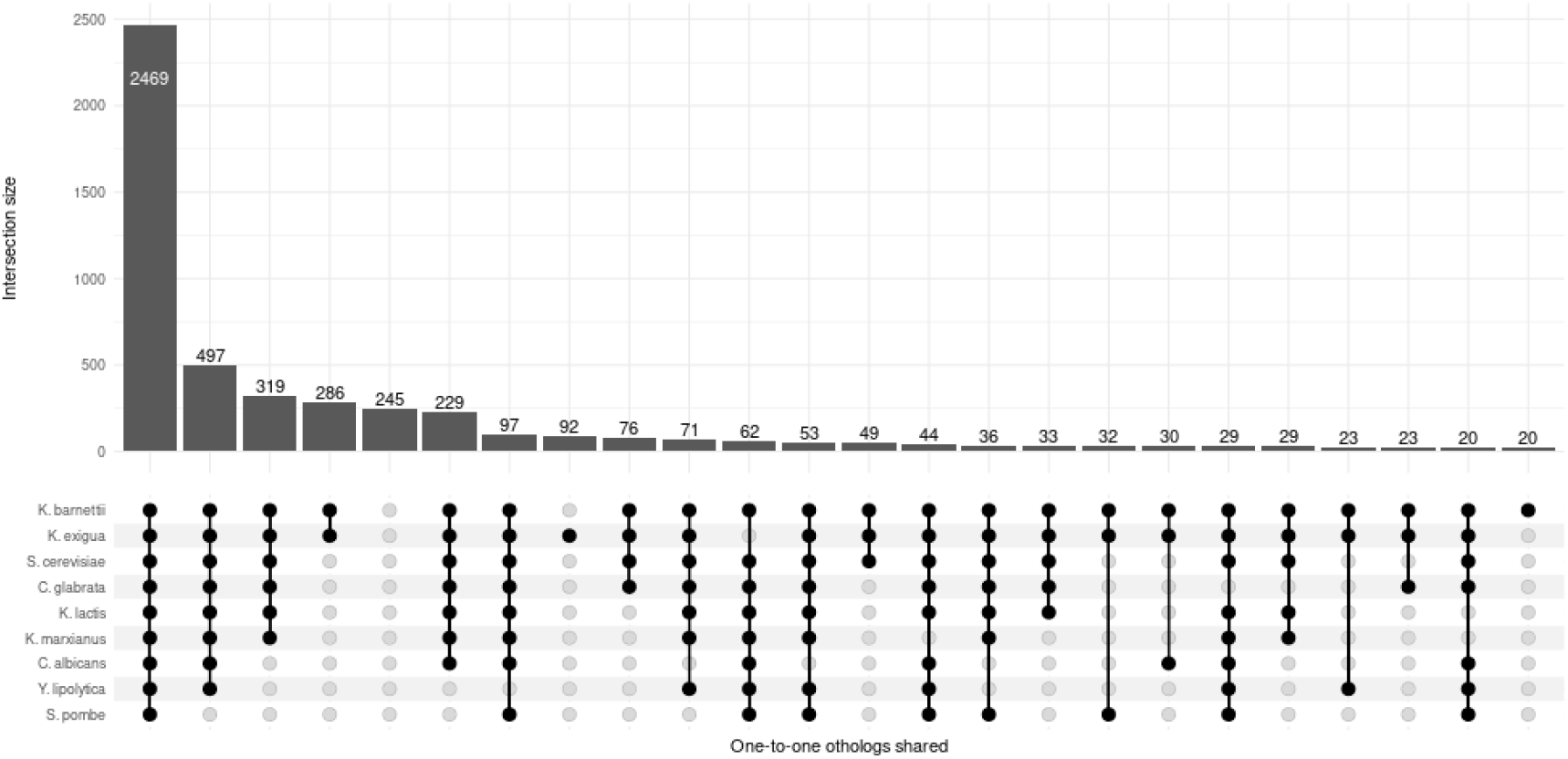
Upset plot showing the number of proteins functionally annotated across *K. bulderi* CBS 8639 strain in common when using *Saccharomyces cerevisiae, Schizosaccharomyces pombe, Candida albicans, Candida glabrata, Yarrowia lipolytica, Kluyveromyces marxianus, Kluyveromyces lactis, Kazachstania exigua* and *Kazachstania barnettii*, as references genomes. Vertical lines and dots across the species represent the proteins in common between the reference species and *K. bulderi*.

Remarkably, 281, 245 and 281 (~5%) proteins of *K. bulderi* CBS 8638, CBS 8639 and NRRL Y-27205 do not have any one-to-one ortholog in any of the yeast species used. 126 genes are shared among the three strains (Dataset S2) and therefore might be *K. bulderi* specific. Interestingly, 193, 160 and 159 out of the potential *K. bulderi* specific proteins, belong to a group of homologs and have been predicted as potential new younger paralogs by HybridMine (Dataset S1).

Algorithms which use the primary sequence homology to assign functional annotation such as HybridMine do not work well on novel sequences. Recent breakthroughs in deep learning are promising for such task. Structure-based protein function prediction using graph convolutional networks have shown to be very accurate in predicting novel protein’s function (26).

We used AlphaFold algorithm to predict the 3D structure of the potential *K. bulderi* specific proteins, and a graph convolutional network model, DeepFRI (27) trained on protein structures and their associated GO terms was used to predict Molecular Function (MF), Biological Process (BP) and Cellular Components (CC) GO terms and EC numbers from the protein structures. Out of the set of potential of *K. bulderi* specific proteins, 42, 40 and 3 had a protein structure predicted by Alphafold in CBS 8638, CBS 8639 and NRRL Y-27205, respectively. Two were the predominant biological processes enriched for these proteins (Dataset S3): RNA related processes and membrane transporters, suggesting that such class of proteins are rapid diverging. Interestingly, it has been shown that across the whole tree of life, cytosolic proteins are under tight selection (*i*.*e*. they are needed for maintaining internal homeostasis), while membrane proteins are under strong adaptive selection. In fact, there are consistently fewer detectable orthologs for membrane proteins than for water-soluble one (28). Moreover, ribosome proteins can present segments of high structural variations which are thought to help adaption to specific environments (29). Therefore, it is not entirely surprisingly that the potential *K. bulderi* specific proteins that have a predicted structure, falls within these two biological processes.

### 4. Structural variation between strains and experimental validation of detected rearrangements

We compared our polished *K. bulderi* genome assemblies by considering CBS 8639 strain as a reference outgroup. We found that all the 12 chromosomes of CBS 8638 are collinear with CBS 8639 (Figure 5A), while NRRL Y-27205 presented a genomic rearrangement in chromosome VII. This region of about 167kb, encompassing 71 genes, was inverted in NRRL Y-27205 genome compared to CBS 8639 (Figure 5B).

**Figure 5.**
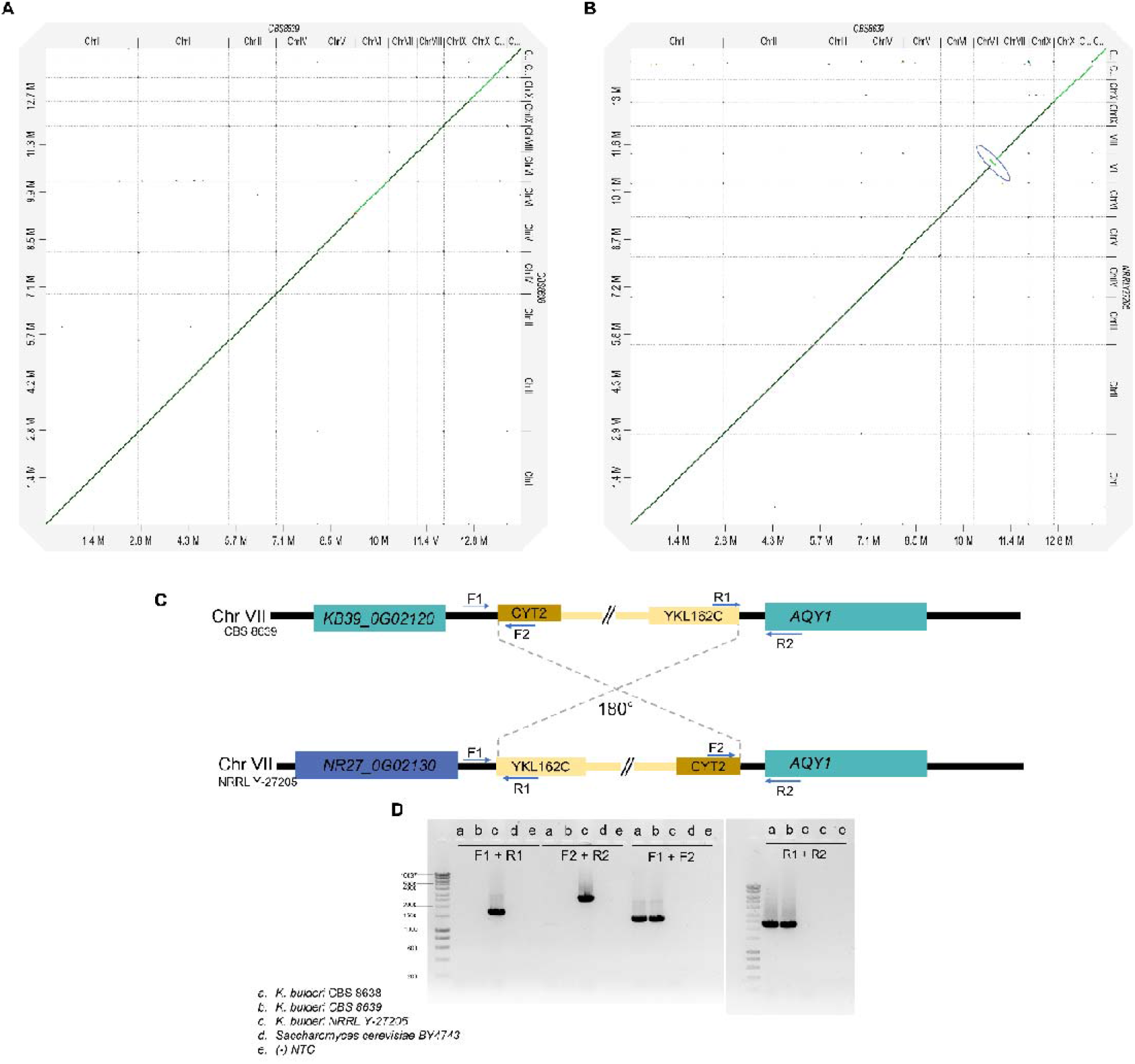
Identification of chromosomal rearrangements between *K. bulderi* strains. Panel A: dot plot representing the alignment between *K. bulderi* CBS 8639 and CBS 8638 genomes; Panel B: dot plot for the alignment between CBS 8639 and NRRL Y-27205 genomes. The inversion in chromosome VII for NRRL Y-27205 is highlighted in blue circle. Panel C: cartoon representation of the inversion including the flanking genes (not to scale) and the location of the primers used to confirm the inversion. Panel D: Gel electrophoresis of PCR products amplified to confirm the inversion. Different combinations of primers (F1-R1, F2-R2, F1-F2 and R1-R2) were used to amplify products in *K. bulderi* CBS 8638 (a), CBS 8639(b) or NRRL Y-27205 (c) and *S. cerevisiae* (d) strains. The negative control (e) has been carried out without DNA as template. In each case the resulting PCR products support the inversion identified by the genome assembly.

We used diagnostic PCR to experimentally confirm this inversion at both sides (Figure 5C). The breakpoints of the inversion in CBS8639 were located in the intergenic regions between *KB39_0G02120* (annotated gene as hypothetical protein) and *KB39_0G02130* (functionally annotated as *CYT2*) on one flank and *KB39_0G02820* (annotated as *AQY1*) and *KB39_0G02810* (annotated as *YKL162C*) on the other flank. The up-stream region in NRRLY-27205 strain does not have the *KB390_G02120* gene, but has different gene, *NR27_0G02130*, in that position, where the breakpoint is. The downstream breakpoint is in the same inter-genic region as CBS 8639. While, *KB39_0G02120* is also an annotated gene in *K. exigua, NR27_0G2130*, is part of the set of genes with no 1:1 ortholog after functional annotated analysis, and no Alphafold prediction.

### 6. Comparison of *K. bulderi* vs other *Kazachstania* genomes

The phylogenetic relationships between *Kazachstania* species remain poorly investigated due to a lack of complete and well-assembled genomes. In fact, only four *Kazachstania* genomes assembled in chromosomes are publicly available, (10–12). *K. barnetti* and *K. saulgeensis* have been isolated in bread sourdough (10,12) and share with *K. bulderi* the ability to growth at low pH, while *K. africana* and *K. naganishii*, isolated in soil (39) and decayed leaves (40) respectively, thus, might not share the same tolerance. The synteny analysis between *K. bulderi* CBS 8639 and these four *Kazachstania* species, revealed an expected high level of conservation with the closest related species, which are also low pH tolerant (Figure 6A). Between *K. bulderi* and its closest relatives *K. barnetti* and *K. saulgensis*, there are 16 and 15 synteny blocks, respectively, larger than 100 Kb and almost all chromosome III is maintained with the same synteny and gene order (Fig. S5 A and B). A breakdown of synteny blocks by their size for all species is reported in Dataset S4 and Figure S5.

**Figure 6.**
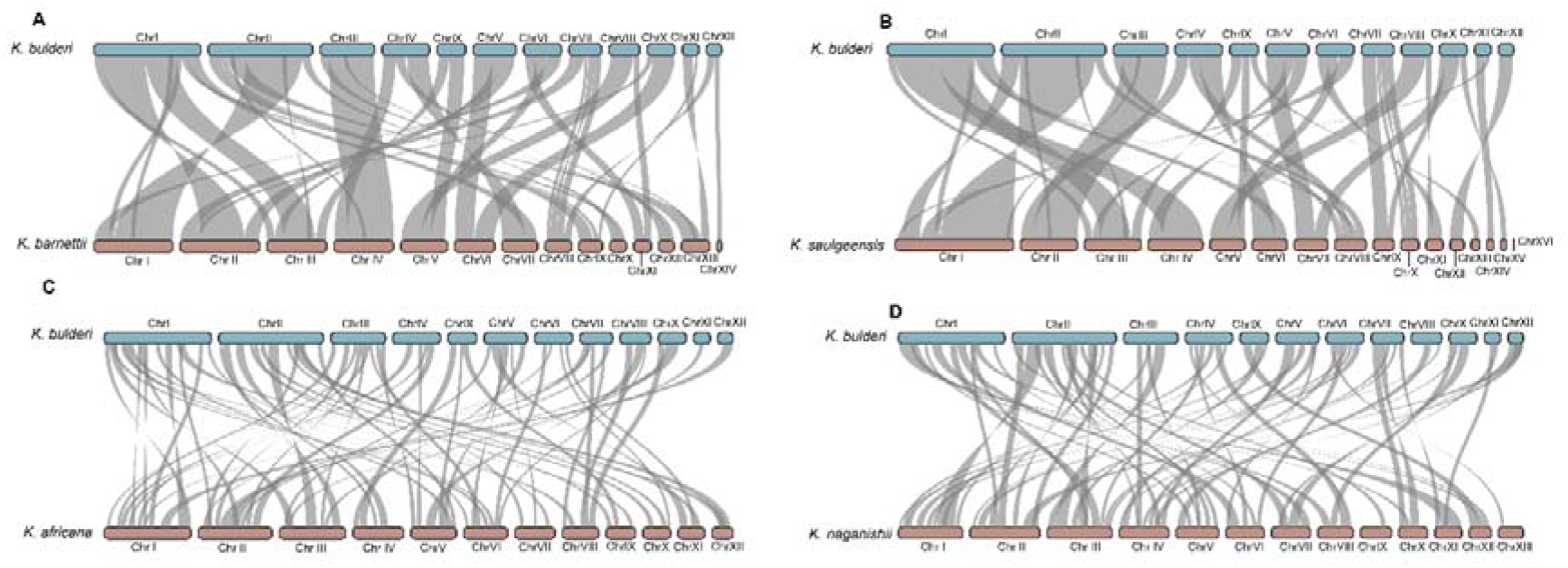
Synteny blocks between the *Kazachstania bulderi* CBS 8639 genome (light blue) and *Kazachstania barnettii* (Panel A), *Kazachstania salugeensis* (Panel B)) *Kazachstania africana* (Panel C*) Kazachstania naganishii* (Panel D). The synteny blocks are represented by the grey areas between the CBS 8639 query genome and the other *Kazachstania sp*. genome.

### 7. Inspection of Mating Type Loci

The *HO* endonuclease gene is present on chromosome I in all strains. The mapping of the MAT locus was carried out using *Saccharomyces cerevisiae, Kazachstania naganishii, Kazachstania saulgeensis* and *Kazachstania barnettii* genomes as reference. We mapped the *HML* and the *MAT* loci in the same region on chromosome V for all three *K. bulderi* strains. With functional annotation we identified only 1:1 ortholog for *MATALPHA1* gene, which however, is located in the HML locus. The HML locus, including the neighbouring genes (*i*.*e. CHA1*) on the X region of the HML locus (Figure 7), is broadly similar to *Kazachstania saulgeensis* and *Kazachstania naganishii* (10). On the Z region of *HML* locus the neighboring genes are different in *K. bulderi* compared with the other *Kazachstania* strains. Such organization of the MAT locus is likely due to a series of deletions following the whole genome duplication (11). There was no mapping for the *HMR* region in the *K. bulderi* strains, suggesting that it might have been lost. The *HMR* locus is conserved between *K. barnettii* and *K. saulgeensis*, (10), but lost in *K. Africana* (11). Moreover, chromosomal rearrangements at the *MAT* locus are common in several *Kazachstania* species. For instance, *K. barnettii* shows an inversion between *HMR* and *MAT* locus (10), whereas in *K. africana* a translocation has caused the loss of *HML* and *HMR* silent cassettes (11).

**Figure 7.**
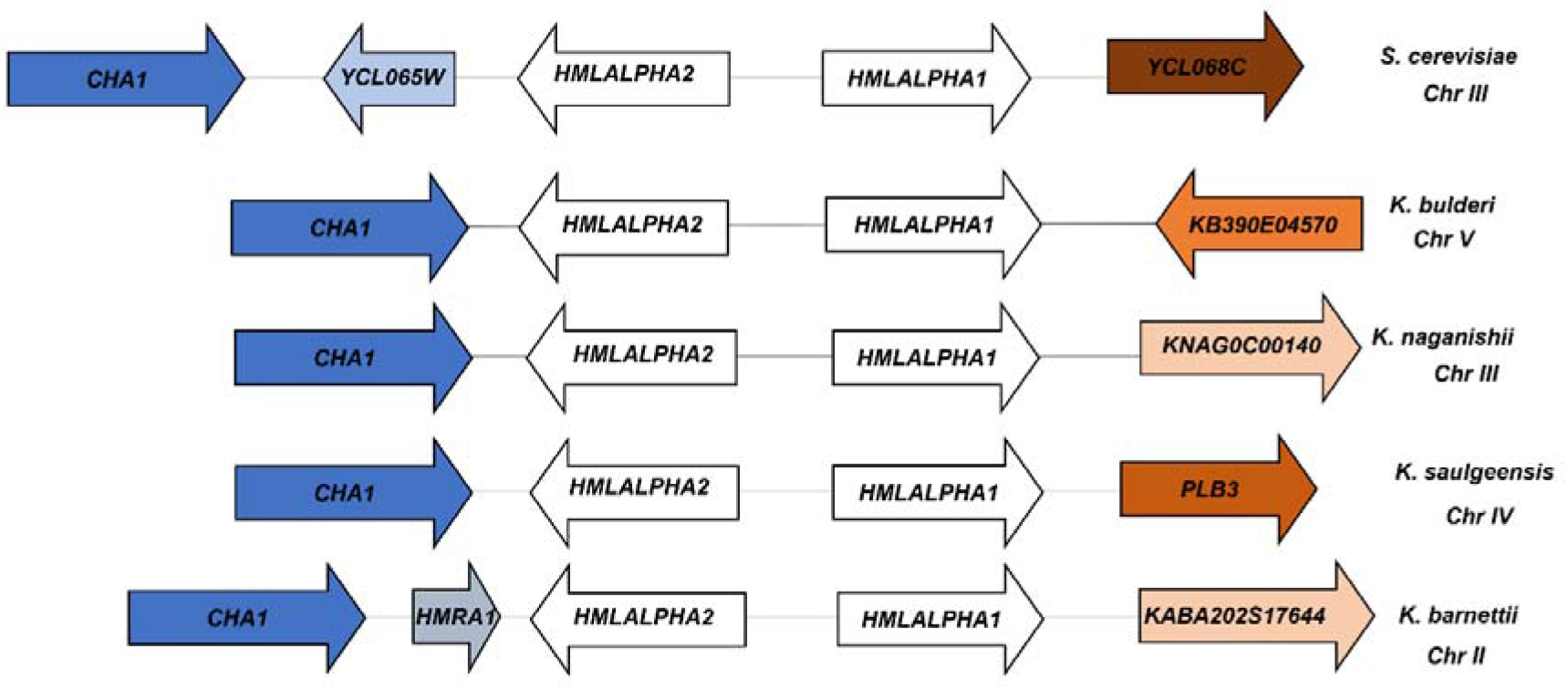
Organization of the HML locus in *K. bulderi, S. cerevisiae, K. naganishii, K. saulgeensis* and *K. barnettii* strains. The blue colored boxes, the central one contains the HML type genes (*i*.*e*., alpha 1 and alpha 2) and the two surrounding ones represent flanking genes. Blocks of the same color between species represent conserved regions with shared syntenic genes.

### 8. Mapping of the mtDNA in *K. bulderi*

Mitochondria have been described to play a crucial role on evolutionary and ecological processes that drive the understanding of phylogenetic relationships between organisms and their mechanisms of adaptation to new conditions (41–43). We could not detect mitochondria DNA using the PacBio suggesting that these strains may be deficient of mitochondrial DNA. This was backed up by the fact that none of them grew on glycerol as sole carbon source (Figure S6). We then use DAPI stain to visualise the DNA content (nuclear and mitochondrial, typically observed in the periphery of the cell) in the three strains, including a rho^+^ and rho^−^ *S. cerevisiae* control strains, which displays a distinct punctate structure (44). We could not detect any mitochondrial DNA for CBS 8638 and CBS 8639 and therefore concluded that these strains are likely to be rho^0^ strains. On the other hand, the staining of the mtDNA of NRRL Y-27205 was similar of the control rho^−^ and contained a mixed population of cells displaying either a rho^−^ or rho^0^ phenotype (Figure 8). To further verify the presence/absence of mitochondrial in these strains, we re-sequenced them using the nanopore technology. We mapped the nanopore reads versus the mtDNA of *Kazatchstania servazzii* of the size of 30,782 bp (15). For CBS 8639 only one read partially mapped to an intergenic region of 200 bp in the mtDNA, while no reads of CBS 8638 mapped to any portion of the *Kazatchstania servazzii* mtDNA, confirming the rho^0^ nature of the strains.

**Figure 8.**
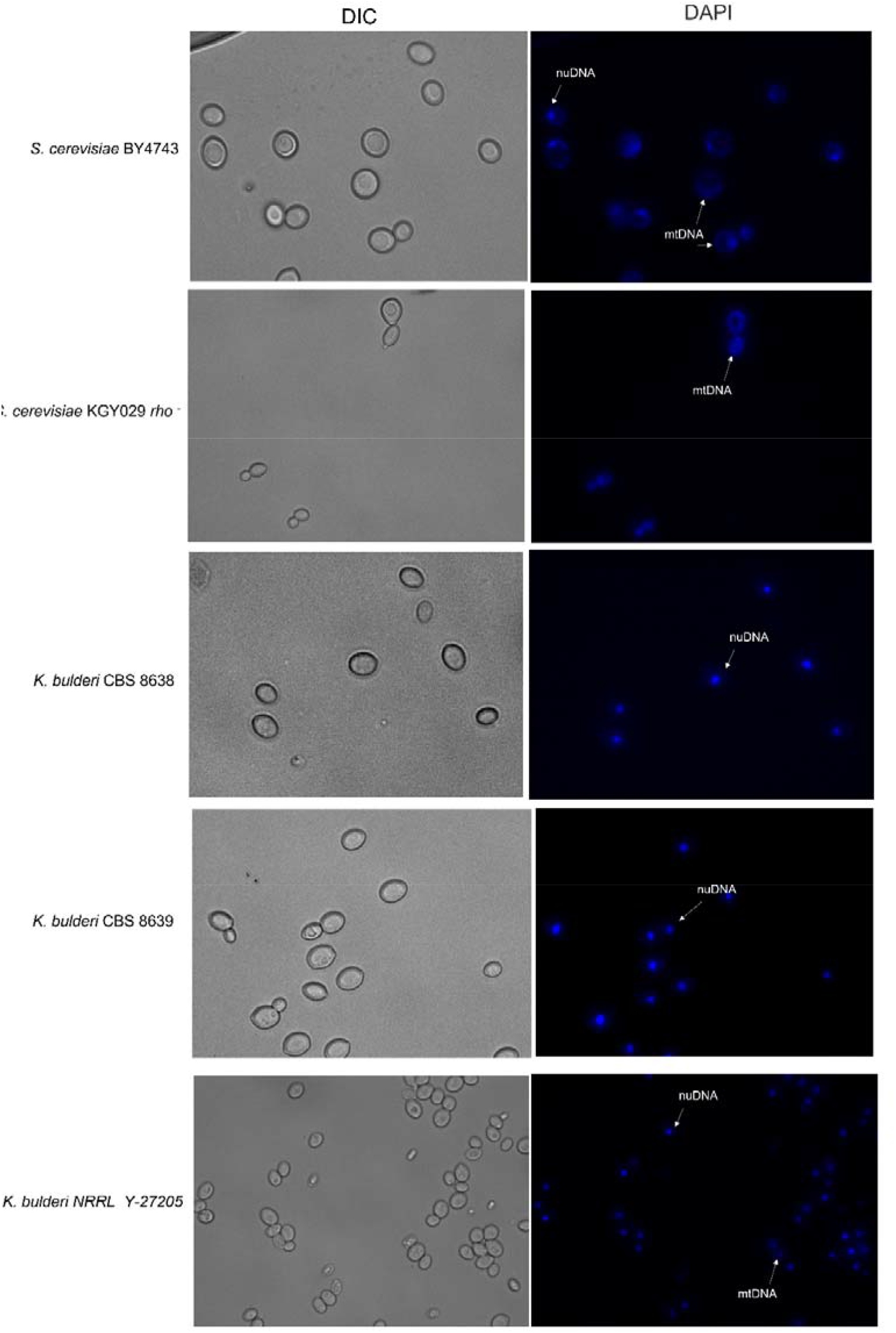
DAPI staining of *K. bulderi* CBS 8638, CBS 8639 and NRRL Y-27205 strains along with *S. cerevisiae* BY4741 and KGY029. Cells were observed under differential interphase contrast (DIC) and after DAPI staining. White arrows point to mitochondrial DNA (mtDNA) and nuclear DNA (nuDNA). The mtDNA in KGY029 shows a network of punctuate dots at the cell periphery, phenotype common in rho^−^ strains.

For NRRL Y-27205 strain we found some evidence of mtDNA, since ca. 2,725 reads mapped to a region of 5,100 bp of the mtDNA from *Kazatchstania servazzii* that includes three genes, namely *COX1, ATP8, ATP6*. We therefore propose to classify NRRL Y-27205 as rho^−^.

## Conclusions

A high-quality genome assembly of three *K. bulderi* strains was achieved through ***i***. a comparison of algorithms for genome assembly and annotation, ***ii***. manual curation and ***iii***. experimental validation and a fully annotated reference genome for this species was constructed. Moreover, we distinguished and validated chromosomal rearrangements in the *K. bulderi* strains studied in this work, that could contribute to their phenotypic variation. Extensive functional annotation revealed potential genus-specific and species-specific genes that might have evolved under the highly selective pressure of maize silage. We showed that the *K. bulderi* strains have low level of heterozygosity and are overall closely related to *K. barnetii* and *K. saulsengensis*, recapitulating the *ITS1* tree. *K. bulderi* strains have lost the mitochondrial DNA and are unable to grow on non-fermentable sources, and they lack the HMR locus.

The current assemblies established a basis for further studying the phylogeny of the *Kazachstania* genus, and it can be used to investigate genomic events associated with yeast domestication and species radiation. Moreover, bespoke molecular tools can now easily be developed for biotechnological purposes, given such strains can optimally grow at low pH and can cope with high concentration of organic acids in the media.

## Materials and Methods

### Strains and phenotypic analysis

Three *K. bulderi* strains CBS 8638, CBS 8639 and NRRL Y-27205 and five *S. cerevisiae* strains 96.2, BY4743, BY4741, NCYC 505 and KGY029 were used in this study. A detailed description of the strains is listed in Table S8.

For liquid fitness assays, *K. bulderi* and *S. cerevisiae* strains were grown at 30°C overnight on YPD and then diluted to an OD_600_ of 0.1. Growth measurements at OD_595_ were recorded in every condition using a FLUOstar OPTIMA Microplate Reader (BMG). The OD_595_ measurements were taken by the microplate reader at 25°C for 72 hours at intervals of 5 min and with 1 min linear shaking before every read. Three technical and three biological samples were used for each *K. bulderi* and *S. cerevisiae* strains. For the lactic acid and the formic acid media, the pH was kept constant to 2.5 and 3, respectively, independently of the acid concentration. To adjust the pH either 10M of NaOH or 1M of H_3_PO_4_ was used.

Graphs were generated by GraphPad Prism, version 7.0 and growth parameters were calculated using “Growthcurver” R package (45). For spot test assays on solid media, cultures were grown overnight at 30°C and number of cells were normalized to an OD_600_ of 4 before being serially diluted 1:10 and spotted onto YPD agar plates containing 100 µg/mL, 150 µg/mL, 200 µg/mL and 300 µg/mL of hygromycin B (Invitrogen), or 5 µg/mL, 10 µg/mL and 25 µg/mL of phleomycin (InvivoGen) and and YP+ 2% glycerol. The drug concentration that inhibits growth in *S. cerevisiae* is 5 µg/mL and 200 µg/mL for phleomycin and hygromycin B, respectively.

### DNA extraction and library preparation for long read next generation sequencing

DNA was extracted from samples using the CTAB (Cetyl trimethyl ammonium bromide) method (46). Briefly, 1mL of overnight culture was added to 50mg of acid washed glass beads 425-600 um (Sigma). 1mL of CTAB extraction buffer was then added, and after incubation at 65°C, RNAse A treatment was conducted by adding 2 uL of RNAse A 100ug/mL (Qiagen) and incubating the samples at 37°C for 15 min. After purification with phenol:chloroform:isoamyl alcohol (25:24:1) the DNA was eluted in 50 uL of ultrapure distilled water (Invitrogen) and store at 4°C. The DNA quality was assessed using the NanoDrop LiTE Spectrophotometer (Thermofisher Scientific) to be within quality specification range required by the PacBio and Oxford Nanopore protocols.

The genomic DNA was adjusted to 10 ng/uL in 150 µL and sheared to approximately 10 kilobase fragments using g-TUBES (Covaris) following the manufacturer’s instructios.. The quality and seize of DNA fragments were verified using Fragment Analyzer (Advanced Analytical Technologies) following the DNF-90 protocol. Samples were prepared for sequencing following the Express Template Prep Kit 2.0 protocol (Pacific Biosciences), with multiplexing using the Barcoded Overhang Adapter kit 8□A (Pacific Biosciences). DNA libraries were sequenced using the SMRT Cell 1□M chips on the Pacific Biosciences Sequel system with 10 hour data acquisition time. For Oxford Nanopore sequencing, 1□µg of the same DNA samples (not sheared) were prepared for sequencing using the SQK-LSK109 Ligation sequencing kit and Flongle sequencing expansion kit FLO-FLG001 (both Oxford Nanopore), following the manufacturer’s instructions. Each strain was sequenced using a MinION Flongle flow cell with 24□hours data acquisition time.

### Genome assembly

Pacbio sequencing data was processed to generate circular consensus sequencing (CCS, or HiFi) reads using the CCS application in SMRT Link 8.0 software package with a minimum three passes, considered to generate a minimum Q20 accuracy. The CCS read length ranged from 10 to 50,000. CCS reads were assembled using the PacBio assemblers algorithms Improved Phased Assembly (IPA v1.8.0) method (available at https://github.com/PacificBiosciences/pbipa.git) and the HiFiasm assembly tool (47) with default settings. Both genome assemblers output consists of one primary contig and one alternate haplotig files that were converted to FASTA format.

### Curation and polishing of the definitive genome assembly

Final adjustments of selected genome assembly were made manually based on the assembly graph, read coverage and distribution and experimental validations. By using *de novo* assemblies of the three *K. bulderi* strains we were able to build a reference genome by ***i***. visualizing the assembly alignment by Bandage V0.8.1 (48); ***ii***. contiguous edges between split contigs (nodes) for each *K. bulderi* strain were identified using the other two strains as references; ***iii***. extracting candidates’ sequences for long PCR validations in order to resolve incomplete/split contigs; ***i****v*. identifying repetitive regions located at the ends of each contigs that are likely to represent telomeric regions; ***v***. mapping the HiFi reads by minimap2 v2.24 (49,50) of each strains against each *de novo* assembly to identify linear read coverage and resolve potential contigs translocated or misplaced by the assembly tool used. The alignments were visualized by Ribbon interactive online visualization tool (51) available http://genomeribbon.com. Primers for long amplicons between contigs edges designed for validations are listed in Supplementary File S1. Finally, assembled sequences were visualized and compared against the final assembly using the BLAST (52), to obtain the final polished assembly. A detailed description of the manual curation is described in Supplementary File S1. Genome assembly statistics about quality and contiguity were assessed using QUAST 5.0.114 (53) at both contig and chromosomal level. To assess completeness we used BUSCO version 5.3.2 (25), based on T=the lineage dataset: saccharomycetes_odb10 (number of genomes: 76, number of BUSCOs: 2137) in both polished and unpolished assemblies. Circos plot of the curated final *K*.*bulderi* genome assembly was generated with Circa (http://omgenomics.com/circa).

### Structural and functional annotations

Gene annotation and gene prediction was achieved using The Yeast Genome Annotation Pipeline (YGAP) (23) and AUGUSTUS (22) for all the *de novo K. bulderi* genome assemblies. The HybridMine tool (24), initially developed for functional annotation at gene level was modified to work at protein level and used to identify one-to-one orthologs between the *K. bulderi* strains and *S. cerevisie, Y. lipolytica, S. pombe, C. albicans, C. glabrata, K. exigua, K. marxianus, K. lactis* and *K. barnetti*, respectively. HybridMine was also used to identify groups of homologs within each *K. bulderi* strain genome. Visualization of the proteins shared between the different species was carried out using “ComplexUpset” and “ggplot2” R packages. Protein structure prediction for 281, 245 and 281 non-annotated proteins in *K. bulderi* CBS 8638, CBS 8639 and NRRL-Y27205, respectively, has been carried out using AlphaFold v2.1.1 (26). Functional annotation from the predicted 3D structures was then achieved using DeepFRI v1.0.0 (27).

### Comparative genome analysis

For analysis of intra and interspecific variation the Burrows–Wheeler Aligner (BWA) software (54) was used to align the sequence reads to the *K. bulderi* newly sequenced genomes. The genomes were first indexed using *bwa index*. The reads were then aligned to their corresponding genome using *bwa aln*. The SAI outputs generated were then converted to SAM format using *bwa samse*. SAMtools suite has been used to process the sequence alignment files (55). The SAM files were converted to BAM files using *samtools view*. BAM files were sorted and indexed for SNP calling using *samtools sort* and *samtools index*, respectively. The SNPs calling was performed using *bcftools mpileup*. Low quality SNPs were filtered using *bcftools view*. The variant analysis was performed inside the CDS regions to calculate heterozygote sites within each genome. The same variant combination and different variant combination between the heterozygote site was also detected. For variances between strains the SNPs and Indels inside and outside CDS were also determined.

The dot plots were generated using D-GENIES (56), an online tool available at http://dgenies.toulouse.inra.fr/. As all *K. bulderi* genome assemblies were constructed *de novo*, alignments were also made to the unorder and initial scaffolds using MUMmer (57) to confirm chromosomal orientation. To determine the level of synteny and visualise the synteny blocks between *K. bulderi* and other *Kazachstania* species the ShinySyn application was used (58–60).

### PCR experimental validation

The continuity of the contigs and the chromosomal rearrangements were validated by PCR. The PCR mixture composed 12.5 μL of 5X LongAmp Taq Reaction Buffer (NEB), 0.4uM of forward and reverse primers (Invitrogen), 2.5U of LongAmp Taq DNA Polymerase (NEB) and 1uL of gDNA. The mix was brought up to 25 μL, final volume, with ultrapure H_2_O. Cycling conditions were set up following manufacturer’s protocol setting the annealing temperature between at 55-58°C. 10 μL of PCR product was loaded on 1% (w/v) agarose gel electrophoresis in 1 x TAE buffer with a 5 μL/100 mL of SafeView nucleic acid stain. Samples were compared to 1 Kb hype ladder (Bioline). The primer sequences are listed in Table S9.

### DAPI staining and mapping of mtDNA

DNA analysis was performed by DAPI (4,6-diamidino-2-phenylindole; Sigma) staining. Yeast cells were harvested after overnight growth in YP+ 2% glycerol and washed twice with PBS. Cells were resuspended in SD media w/o amino acids. DAPI and SDS were added to the culture at the final concentration of 1 µg/ml and 0.01%, respectively. The cells were incubated in the dark for 10 min at 30°C. Cells were observed with a Eclipse TE2000-U fluorescence inverted microscope (Nikon) fitted with a × 100 immersion objective. The images were capture using the Ocular Image Acquisition Software V2.0 (QImaging). The images were then processed and assembled with Image J (61,62).

For Oxford Nanopore data, base calling was performed using Guppy (Oxford Nanopore) and the reads from CBS 8638, CBS 8639 and NRRL Y-27205 were aligned against the sequence of mtDNA from *K. servazzii* using Minimap2 v2.24. The aligned mitochondria reads were extracted and remapped against the *K. bulderi* genome assemblies to rule out mapping to previous annotated nuclear genes.

## Acknowledgments

This work was supported by BBSRC-link grant (BB/T002123/1) awarded to DD, and we want to acknowledge the economic support provided by BP for this project. AH is supported by a studentship from the Future Biomanufacturing hub and BP. We thank Dr. Ling Li for the valuable discussions and support at the beginning of the project, and Dr. Kirk Malone for commercial insights. We also thank Marco Monti and Dr. Kewin Gombeau for the support on genomic analysis tools, and for providing the KGY029 strain and guidance for mtDNA analysis, respectively.

## References

1. Tejayadi S, Cheryan M. Lactic acid from cheese whey permeate. Productivity and economics of a continuous membrane bioreactor. Appl Microbiol Biotechnol 1995 432. 1995;43(2):242–8.

2. Ahmad A, Banat F, Taher H. A review on the lactic acid fermentation from low-cost renewable materials: Recent developments and challenges. Environ Technol Innov. 2020 Nov 1;20:101138.

3. Prado-Rubio OA, Gasca-González R, Fontalvo J, Gómez-Castro FI, Pérez-Cisneros ES, Morales-Rodriguez R. Design and evaluation of intensified downstream technologies towards feasible lactic acid bioproduction. Chem Eng Process - Process Intensif. 2020 Dec 1;158:108174.

4. Lee HD, Lee MY, Hwang YS, Cho YH, Kim HW, Park HB. Separation and Purification of Lactic Acid from Fermentation Broth Using Membrane-Integrated Separation Processes. Ind Eng Chem Res. 2017 Jul 26;56(29):8301–10.

5. Baral P, Pundir A, Kurmi A, Singh R, Kumar V, Agrawal D. Salting-out assisted solvent extraction of L (+) lactic acid obtained after fermentation of sugarcane bagasse hydrolysate. Sep Purif Technol. 2021 Aug 15;269:118788.

6. Chen Y, Nielsen J. Biobased organic acids production by metabolically engineered microorganisms. Curr Opin Biotechnol. 2016 Feb 1;37:165–72.

7. Middelhoven WJ, Kurtzman CP, Vaughan-Martini A. Saccharomyces bulderi sp. nov., a yeast that ferments gluconolactone. Antonie van Leeuwenhoek 2000 773. 2000;77(3):223–8.

8. Dijken JP van, Tuijl A van, Luttik MAH, Middelhoven WJ, Pronk JT. Novel Pathway for Alcoholic Fermentation of δ-Gluconolactone in the Yeast Saccharomyces bulderi. J Bacteriol. 2002;184(3):672.

9. Kurtzman Cletus P. Fell Jack W. Boekhout T. The yeasts. 5th ed. Elsevier; 2011. 439–470 p.

10. Devillers H, Sarilar V, Grondin C, Sterck L, Segond D, Jacques N, et al. Whole-Genome Sequences of Two Kazachstania barnettii Strains Isolated from Anthropic Environments. Genome Biol Evol. 2022 Feb 4;14(2).

11. Gordon JL, Armisen D, Proux-Weŕa E, Oh Eígeartaigh SS, Byrne KP, Wolfe KH. Evolutionary erosion of yeast sex chromosomes by mating-type switching accidents. Proc Natl Acad Sci U S A. 2011 Dec 13;108(50):20024–9.

12. Sarilar V, Sterck L, Matsumoto S, Jacques N, Neuvéglise C, Tinsley CR, et al. Genome sequence of the type strain CLIB 1764T (= CBS 14374T) of the yeast species Kazachstania saulgeensis isolated from French organic sourdough. Genomics Data. 2017 Sep 1;13:41.

13. García-Ortega LF, Colón-González M, Sedeño I, Santiago-Garduño E, Avelar-Rivas JA, Kirchmayr MR, et al. Draft Genome Sequence of a Kazachstania humilis Strain Isolated from Agave Fermentation. Microbiol Resour Announc. 2022 Mar 17;11(3).

14. Faherty L, Lewis C, McElheron M, Garvey N, Duggan R, Shovlin B, et al. Draft Genome Sequences of Two Isolates of the Yeast Kazachstania servazzii Recovered from Soil in Ireland. Microbiol Resour Announc. 2019 Oct 31;8(44).

15. Langkjær RB, Casaregola S, Ussery DW, Gaillardin C, Piškur J. Sequence analysis of three mitochondrial DNA molecules reveals interesting differences among Saccharomyces yeasts. Nucleic Acids Res. 2003 Jun 15;31(12):3081–91.

16. Morio F, O’Brien CE, Butler G. Draft Genome Sequence of the Yeast Kazachstania telluris CBS 16338 Isolated from Forest Soil in Ireland. Mycopathol 2020 1853. 2020 Apr 30;185(3):587–90.

17. Davies CP, Arfken AM, Frey JF, Summers KL. Draft Genome Sequence of Kazachstania slooffiae, Isolated from Postweaning Piglet Feces. Microbiol Resour Announc. 2021 Aug 26;10(34).

18. Deroche L, Buyck J, Cateau E, Rammaert B, Marchand S, Brunet K. Draft Genome Sequence of Kazachstania bovina Yeast Isolated from Human Infection. Mycopathologia. 2022 Aug 1;187(4):413–5.

19. Kaeuffer C, Baldacini M, Ruge T, Ruch Y, Zhu YJ, De Cian M, et al. Fungal Infections Caused by Kazachstania spp., Strasbourg, France, 2007–2020. Emerg Infect Dis. 2022 Jan 1;28(1):30.

20. Shen XX, Opulente DA, Kominek J, Zhou X, Steenwyk JL, Buh K V., et al. Tempo and Mode of Genome Evolution in the Budding Yeast Subphylum. Cell. 2018 Nov 29;175(6):1533–1545.e20.

21. Molina-Mora JA, Campos-Sánchez R, Rodríguez C, Shi L, García F. High quality 3C de novo assembly and annotation of a multidrug resistant ST-111 Pseudomonas aeruginosa genome: Benchmark of hybrid and non-hybrid assemblers. Sci Reports 2020 101. 2020 Jan 29;10(1):1–16.

22. Stanke M, Morgenstern B. AUGUSTUS: a web server for gene prediction in eukaryotes that allows user-defined constraints. Nucleic Acids Res. 2005 Jul 7;33(Web Server issue):W465.

23. Proux-Wéra E, Armisén D, Byrne KP, Wolfe KH. A pipeline for automated annotation of yeast genome sequences by a conserved-synteny approach. BMC Bioinformatics. 2012 Sep 17;13(1):1–12.

24. Timouma S, Schwartz JM, Delneri D. HybridMine: A Pipeline for Allele Inheritance and Gene Copy Number Prediction in Hybrid Genomes and Its Application to Industrial Yeastss. Microorganisms. 2020 Oct 1;8(10):1–15.

25. Manni M, Berkeley MR, Seppey M, Simão FA, Zdobnov EM. BUSCO Update: Novel and Streamlined Workflows along with Broader and Deeper Phylogenetic Coverage for Scoring of Eukaryotic, Prokaryotic, and Viral Genomes. Mol Biol Evol. 2021 Sep 27;38(10):4647–54.

26. Jumper J, Evans R, Pritzel A, Green T, Figurnov M, Ronneberger O, et al. Highly accurate protein structure prediction with AlphaFold. Nat 2021 5967873. 2021 Jul 15;596(7873):583–9.

27. Gligorijević V, Renfrew PD, Kosciolek T, Leman JK, Berenberg D, Vatanen T, et al. Structure-based protein function prediction using graph convolutional networks. Nat Commun. 2021 Dec 1;12(1).

28. Sojo V, Dessimoz C, Pomiankowski A, Lane N. Membrane Proteins Are Dramatically Less Conserved than Water-Soluble Proteins across the Tree of Life. Mol Biol Evol. 2016 Nov 1;33(11):2874.

29. Melnikov S, Manakongtreecheep K, Söll D. Revising the Structural Diversity of Ribosomal Proteins Across the Three Domains of Life. Mol Biol Evol. 2018 Jul 1;35(7):1588.

30. Liti G, Nguyen Ba AN, Blythe M, Müller CA, Bergström A, Cubillos FA, et al. High quality de novo sequencing and assembly of the Saccharomyces arboricolus genome. BMC Genomics. 2013 Jan 31;14(1):1–14.

31. Peter J, De Chiara M, Friedrich A, Yue JX, Pflieger D, Bergström A, et al. Genome evolution across 1,011 Saccharomyces cerevisiae isolates. Nat 2018 5567701. 2018 Apr 11;556(7701):339–44.

32. Plech M, de Visser JAGM, Korona R. Heterosis is prevalent among domesticated but not wild strains of Saccharomyces cerevisiae. G3 Genes, Genomes, Genet. 2014 Feb 1;4(2):315–23.

33. Muller LAH, McCusker JH. Microsatellite analysis of genetic diversity among clinical and nonclinical Saccharomyces cerevisiae isolates suggests heterozygote advantage in clinical environments. Mol Ecol. 2009 Jul;18(13):2779.

34. Rodrigues-Prause A, Sampaio NMV, Gurol TM, Aguirre GM, Sedam HNC, Chapman MJ, et al. A Case Study of Genomic Instability in an Industrial Strain of Saccharomyces cerevisiae. G3 Genes|Genomes|Genetics. 2018 Nov 1;8(11):3703.

35. Beekman CN, Ene I V. Short-term evolution strategies for host adaptation and drug escape in human fungal pathogens. PLOS Pathog. 2020 May 1;16(5):e1008519.

36. Forche A, Abbey D, Pisithkul T, Weinzierl MA, Ringstrom T, Bruck D, et al. Stress alters rates and types of loss of heterozygosity in candida albicans. MBio. 2011;2(4).

37. Lancaster SM, Payen C, Heil CS, Dunham MJ. Fitness benefits of loss of heterozygosity in Saccharomyces hybrids. Genome Res. 2019 Oct 1;29(10):1685–92.

38. Sui Y, Qi L, Wu JK, Wen XP, Tang XX, Ma ZJ, et al. Genome-wide mapping of spontaneous genetic alterations in diploid yeast cells. Proc Natl Acad Sci U S A. 2020 Nov 10;117(45):28191–200.

39. Van der Walt JP. The yeast Kluyveromyces africanus nov. spec. and its phylogenetic significance. Antonie Van Leeuwenhoek. 1956 Dec;22(4):321–6.

40. Mikata K, Ueda-Nishimura K, Hisatomi T. Three new species of Saccharomyces sensu lato van der Walt from Yaku Island in Japan: Saccharomyces naganishii sp. nov., Saccharomyces humaticus sp. nov. and Saccharomyces yakushimaensis sp. nov. Int J Syst Evol Microbiol. 2001;51(Pt 6):2189–98.

41. Hill GE. Mitonuclear coevolution as the genesis of speciation and the mitochondrial DNA barcode gap. Ecol Evol. 2016 Aug 1;6(16):5831–42.

42. De Chiara M, Friedrich A, Barré B, Breitenbach M, Schacherer J, Liti G. Discordant evolution of mitochondrial and nuclear yeast genomes at population level. BMC Biol 2020 181. 2020 May 11;18(1):1–15.

43. Gershoni M, Templeton AR, Mishmar D. Mitochondrial bioenergetics as a major motive force of speciation. BioEssays. 2009 Jun 1;31(6):642–50.

44. Okamoto S, Inai T, Miyakawa I. Morphology of mitochondrial nucleoids in respiratory-deficient yeast cells varies depending on the unit length of the mitochondrial DNA sequence. FEMS Yeast Res. 2016 Aug 1;16(5):55.

45. Sprouffske K, Wagner A. Growthcurver: An R package for obtaining interpretable metrics from microbial growth curves. BMC Bioinformatics. 2016 Apr 19;17(1):1–4.

46. Wu ZH, Wang TH, Huang W, Qu YB. A simplified method for chromosome DNA preparation from filamentous fungi. Mycosystema. 2001;20(4):575.

47. Cheng H, Concepcion GT, Feng X, Zhang H, Li H. Haplotype-resolved de novo assembly using phased assembly graphs with hifiasm. Nat Methods 2021 182. 2021 Feb 1;18(2):170–5.

48. Wick RR, Schultz MB, Zobel J, Holt KE. Bandage: interactive visualization of de novo genome assemblies. Bioinformatics. 2015 Oct 15;31(20):3350–2.

49. Li H. New strategies to improve minimap2 alignment accuracy. Bioinformatics. 2021;37(23):4572–4.

50. Li H. Minimap2: pairwise alignment for nucleotide sequences. Bioinformatics. 2018;34(18):3094–100.

51. Nattestad M, Aboukhalil R, Chin CS, Schatz MC. Ribbon: intuitive visualization for complex genomic variation. Bioinformatics. 2021 Apr 20;37(3):413–5.

52. Altschul SF, Gish W, Miller W, Myers EW, Lipman DJ. Basic local alignment search tool. J Mol Biol. 1990 Oct 5;215(3):403–10.

53. Gurevich A, Saveliev V, Vyahhi N, Tesler G. QUAST: quality assessment tool for genome assemblies. Bioinformatics. 2013 Apr 15;29(8):1072–5.

54. Li H, Durbin R. Fast and accurate short read alignment with Burrows–Wheeler transform. Bioinformatics. 2009 Jul 15;25(14):1754–60.

55. Danecek P, Bonfield JK, Liddle J, Marshall J, Ohan V, Pollard MO, et al. Twelve years of SAMtools and BCFtools. Gigascience. 2021 Jan 29;10(2).

56. Cabanettes F, Klopp C. D-GENIES: Dot plot large genomes in an interactive, efficient and simple way. PeerJ. 2018 Jun 4;2018(6):e4958.

57. Kurtz S, Phillippy A, Delcher AL, Smoot M, Shumway M, Antonescu C, et al. Versatile and open software for comparing large genomes. Genome Biol. 2004 Jan 30;5(2):1–9.

58. Kiełbasa SM, Wan R, Sato K, Horton P, Frith MC. Adaptive seeds tame genomic sequence comparison. Genome Res. 2011 Mar;21(3):487–93.

59. Tang H, Bowers JE, Wang X, Ming R, Alam M, Paterson AH. Synteny and Collinearity in Plant Genomes. Science (80-). 2008 Apr 25;320(5875):486–8.

60. Xiao Z, Lam H-M. ShinySyn: a Shiny/R application for the interactive visualization and integration of macro- and micro-synteny data. Bioinformatics. 2022 Sep 15;38(18):4406–8.

61. Abràmoff MD, Magalhães PJ, Ram SJ. Image processing with ImageJ. Biophotonics Int. 2004;11(7):36–42.

62. Schneider CA, Rasband WS, Eliceiri KW. NIH Image to ImageJ: 25 years of image analysis. Nat Methods 2012 97. 2012 Jun 28;9(7):671–5.

