## Supplementary material for "High quality *de novo* genome assembly of non-conventional yeast *Kazachstania bulderi* a new potential low pH production host for biorefineries": Supplementary File 1 Manual Curation.docx

**Supplementary File: Manual curation**

***i.* Continuity of contigs**

To resolve incomplete/split contigs in the IPA assemblies deep comparison between the initial assemblies of the three *K. bulderi* were carried out by aligning CBS 8639 genome assembly against CBS 8638 and NRRL Y-27205. All *de novo* assembly graphs were visualized using Bandage v0.8.1 (1) and the alignment showed single contigs that have been assembled separated on two or even three different contigs (Table SF1). For instance, contig ctg.00005F of CBS8639 was split in contig ctg.00006F, ctg.00026F, ctg.00027F in CBS 8638 initial assembly (Figure SF1).

Candidate sequences for long PCR validation were extracted by selecting the regions at the edge of each the split/incomplete contig. We validated the continuity of chromosome 1, 3, 4, 8, 10 and 12 that were broken in two different contigs and also chromosome 7 that was split in three different contigs in CBS 8638 assembly (Figure SF1C). A list of primers with a detailed explanation of potential split contigs and amplicon sizes are listed in Table SF2.

The continuity of the contigs were further verified by Mapping the CCS reads of each strain against each de novo assembly by Minimap2 v2.24 (2,3). We identified lineal read coverage and resolve potential contigs translocated or misplaced by the IPA assembler by visualizing the alignment using Ribbon (4).

**Table SF1**. Summary of all the connections of the initial primary contigs. Similar contigs were assign the same number followed by letters a, b, c to show connections.

| Strain | Name Original assembly IPA | Initial nomenclature |
| --- | --- | --- |
| CBS8639 | ctg.000000F | LB Chr I |
|  | ctg.000001F | LB Chr II |
|  | ctg.000006F | LB Chr III |
|  | ctg.000008F | LB Chr IV |
|  | ctg.000003F | LB Chr V |
|  | ctg.000011F | LB Chr VI |
|  | ctg.000005F | LB Chr VII |
|  | ctg.000016F | LB Chr VIII a |
|  | ctg.000023F | LB Chr VIII b |
|  | ctg.000009F | LB Chr IX |
|  | ctg.000015F | LB Chr X |
|  | ctg.000020F | LB Chr XI |
|  | ctg.000014F | LB Chr XII |
|  | ctg.000022F | LB Chr XIII |
| CBS8638 | ctg.000001F | LB Chr I a |
|  | ctg.000002F | LB Chr I b |
|  | ctg.000015F | LB Chr I c |
|  | ctg.000000F | LB Chr II |
|  | ctg.000005F | LB Chr III |
|  | ctg.000004F | LB Chr IV |
|  | ctg.000008F | LB Chr V |
|  | ctg.000012F | LB Chr VI |
|  | ctg.000027F | LB Chr VII a |
|  | ctg.000026F | LB Chr VII b |
|  | ctg.000006F | LB Chr VII c |
|  | ctg.000014F | LB Chr VIII |
|  | ctg.000007F | LB Chr IX |
|  | ctg.000018F | LB Chr X a |
|  | ctg.000017F | LB Chr X b |
|  | ctg.000020F | LB Chr XI |
|  | ctg.000019F | LB Chr XII a |
|  | ctg.000015F | LB Chr XII b |
|  | ctg.000004F | LB Chr XIII |
| NRRL Y-27205 | ctg.000000F | LB Chr I |
|  | ctg.000001F | LB Chr II |
|  | ctg.000010F | LB Chr III a |
|  | ctg.000020F | LB Chr III b |
|  | ctg.000008F | LB Chr IV a |
|  | ctg.000024F | LB Chr IV b |
|  | ctg.000003F | LB Chr V |
|  | ctg.000009F | LB Chr VI |
|  | ctg.000005F | LB Chr VII |
|  | ctg.000013F | LB Chr VIII |
|  | ctg.000002F | LB Chr IX |
|  | ctg.000015F | LB Chr X |
|  | ctg.000017F | LB Chr XI |
|  | ctg.000018F | LB Chr XII |
|  | ctg.000023F | LB Chr XIII |

**Table SF2.** Primers for experimental validation of split/incomplete contigs for CBS 8638, CBS8639 and NRRL Y-27205 IPA genome assemblies.

| ctg.000016F and ctg000023F of CBS 8639 form Chr 8 | | |
| --- | --- | --- |
| **Chr 8 F** | GGAACTTCCCGCATCATTTT | |
| **Chr 8 R** | GCTGCTGAGGCTAATGAATT | |
| Amplicon size | 1129 | (bp) |
| ctg.000010F and ctg.000020F in NRRL Y-27205 form Chr 3 | | |
| **Chr 3 F** | GTCTAGCACGTAACCTTGGT | |
| **Chr 3 R** | CGATAATGATTCGGTTGTTCC | |
| Amplicon size | 1300 | (bp) |
| ctg.00008F and ctg.000024F verification for Chr IV | | |
| **Chr 4 F** | CTAGGTAATTGAGCAGGTGG | |
| **Chr 4 R** | CGGTTACACTACTAACGACC | |
| Amplicon size | 1691 | (bp) |
| **Chr 7 splits in three contigs** | |  |
| Starts in ctg. 000027F follow by ctg.000026F finishes in ctg.000006F | | |
| Ch7A F | CCAGCTTCGATATCTTCAGC | |
| Ch7A R | ACCAACCGTTCGAGTATATC | |
| Amplicon size | 1304 | (bp) |
| Ch7B F | AAGGCTGAACCAACTCCGGA | |
| Ch7B R | TAGCACGGATTTAGTGTCCCG | |
| Amplicon size | 1107 | (bp) |
| **Chr 1a + b** |  |  |
| Start ctg.00001F and ctg.000002F in CBS 8638 | |  |
| Chr 1A F | GGTACCCTCAGGGTATTCTG | |
| Chr 1A R | CCGCAGGTTCTAATCAAGCT | |
| Amplicon | 800 | (bp) |
| **Chr XII** |  |  |
| ctg.000019F and ctg.000015F | |  |
| Chr 12 F | GGAGCGTAACATCACAGACC | |
| Chr 12 R | GCCATTAGACCTGGCATTTC | |
| Amplicon size | 834 | (bp) |
| **Chr X** |  |  |
| ctg.000018F and cgt.000017F form chr X | |  |
| Chr 10 F | GGTTGCACTGACTTGAGTTC | |
| Chr 10 R | CGGATGGAATATCCGATGATG | |
| Amplicon size | 789 | (bp) |

**Figure SF1**

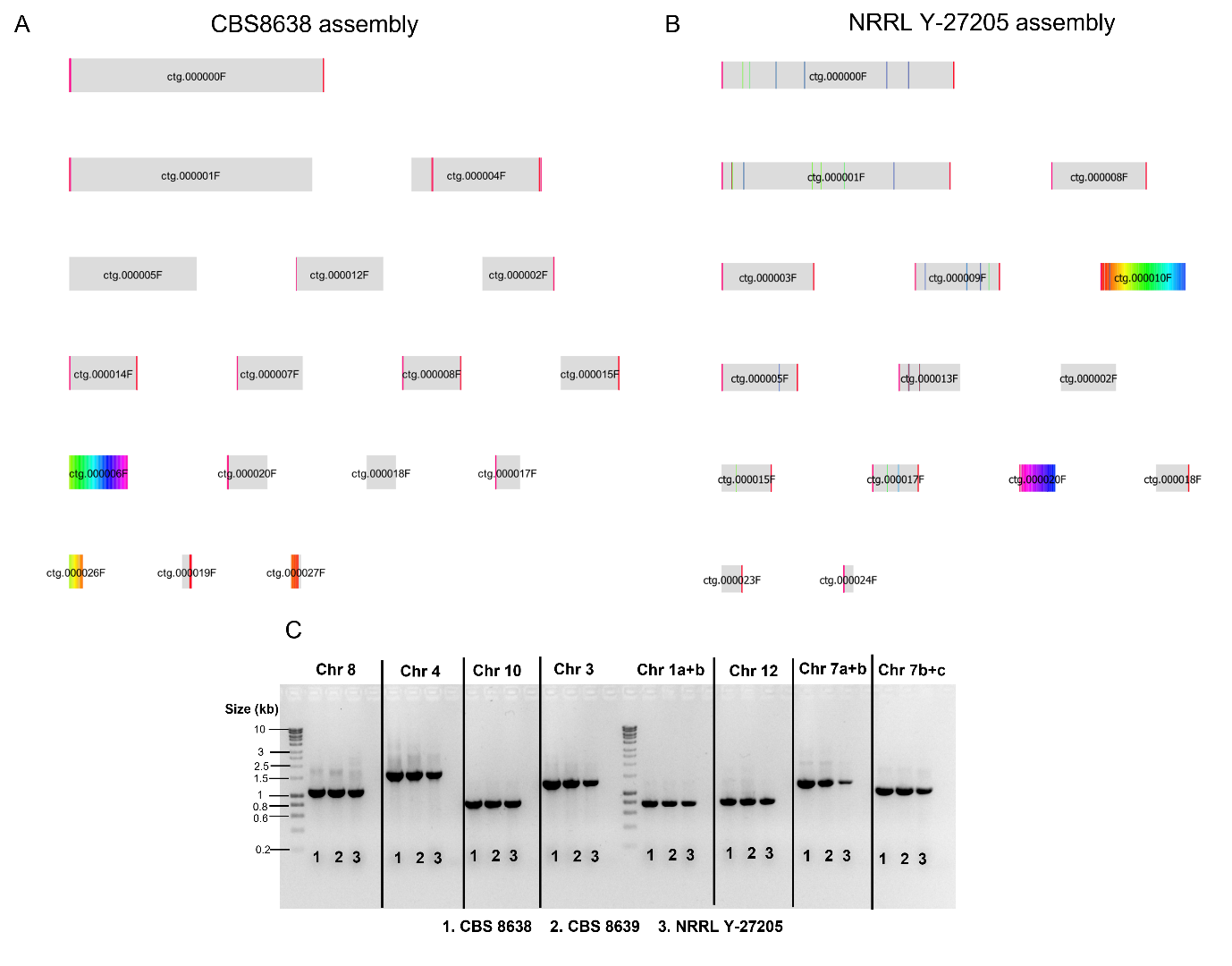

**Figure SF1**. Panel A: representation of initial CBS8638 assembly (17 contigs) blasted against CBS8639 (14 contigs) initial assembly; Panel B: NRRL Y-27205 assembly (15 contigs) blasted against CBS8639. Rainbow colours (from red to violet) represents the continuity of the contigs in CBS8639 with the best BLAST hits. For instance, contig ctg.00005F of CBS8639 is divided in ctg.000027F, followed by ctg.000026F and finishing in ctg.00006F for CBS8638. In Panel B contig ctg. 00006F in CBS8639 is divided in contig ctg. 000010F and ctg.000020F in NRRL Y-27205. Panel C: PCR verification of contigs breakpoints for CBS8638, CBS8639 and NRRL Y-27205 strains for chromosomes 1, 3, 4, 7, 8, 10, and 12.

***ii.* Assembly of Chromosome I and V**

By mapping CBS 8638 and CBS 8639 reads against CBS 8638 assembly, we identified non-continuous coverage in the named chromosome I and chromosome V. A region at the end of chromosome I reads were interrupted for reads that were starting on chromosome V and continuing on chromosome I (Figure SF2A), pattern similar to the one observed on translocations. To investigate whether there is a translocation or a miss-assembly, we experimentally amplify by different regions of chromosome I and chromosome V by PCR the using primers located up-stream and downstream of coverage breakpoint (Figure SF2B). A list of the primers used for this PCR validation is shown in Table SF3. There was no amplification for the possible translocated region, while bands for the primers located on the same chromosome showed continuity for both, chromosome V and I. This result rules out the translocation and proved that assembly algorithms are prompt to mis-assemblies, and manual curation is a crucial step for a high-quality *de novo* genome assembly. After correcting the assembly and aligned the reads to the polished CBS 8638 assembly continuous coverage was achieved (Figure SF2C).

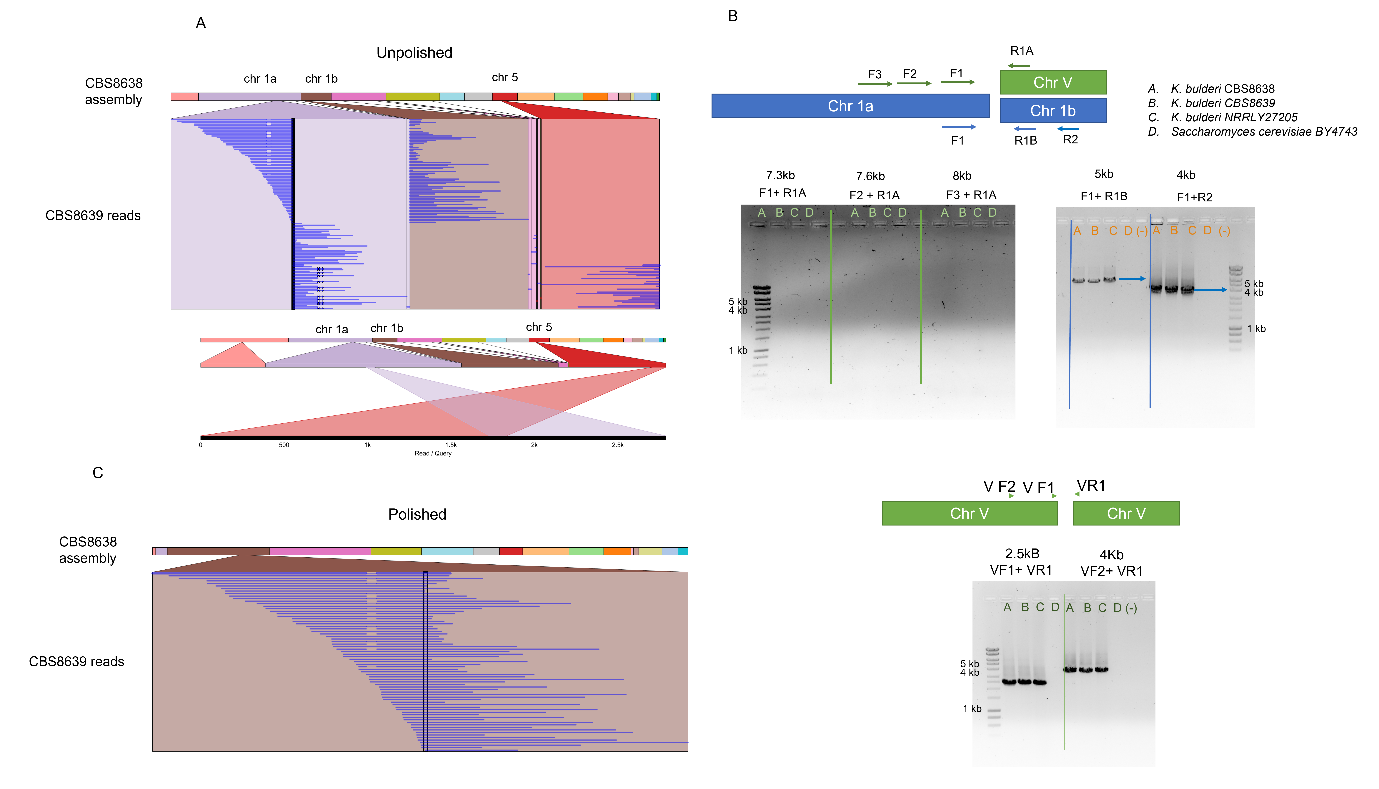

**Figure SF2.** Panel A: visual representation of the alignment of CBS 8639 reads in the region of chromosome I that was identify as potentially translocated in chromosome V in the unpolished *K. bulderi* CBS 8638 genome assembly. Panel B: representation of the location of primers for PCR verification of the continuity of chromosome I and chromosome V and PCR-based confirmation of the continuity of chr I and V. Different primer combinations were used to amplify products corresponding to either CBS 8638, CBS 8639 or NRRL Y-27205 strain. *S. cerevisiae* BY4743 was used as negative control. In each case the resulting PCR products support the miss-assembly identified and ruled out the translocation. Panel C: alignment of CBS 8639 reads against CBS 8638 genome assembly after curation.

**Table SF3**

| **Name** | **Sequence** |
| --- | --- |
| F1 | CATCCTTGGAGACGTTAGCTC |
| R1 A | GACCACACTCCAAACGGTCT |
| *Expected band* | 8821 |
| F2 | GGAGAGAATCTCGCGGTGAG |
| *Expected band* | 7658 |
| F3 | GGAGTGTCGATAGCCATTCTAAC |
| *Expected band* | 7305 |
| F1' | CGGCTAATGAAATCCTTCCTC |
| R1B | CTCTACTGCTAAGCCAGGTG |
| *Expected band* | 5901 |
| R2 | GCAGTGATGGCTCAAATGTC |
| *Expected band* | 4081 |
| VF1 | CCAACCACAACGTGTAGTG |
| VR1 | GACCACACTCCAAACGGTCT |
| *Expected band* | 2556 |
| VF2 | GGCACTTGAGACATATGAGC |
| *Expected band* | 4026 |

***iii.* Missing coverage**

About 200,000 bp coverage was missing from chromosome V (Figure SF3A). This region was present on CBS 8638 but was not present on CBS 8639 and NRRL Y-27205. To identify whether the section of DNA was present on all strains or not, we used the alternative haploitgs to find the missing sequence. The missing region was also validated by PCR and the reads were re-mapped to verify continuity (Figure SF3B). We recovered around 56 genes located in the missing region.

The chromosome like polished assembly reduced the mismatches per 100kb from 706.3 to 695.96 in CBS 8638 and from 842.98 to 824.63 in NRRL Y-27205.

**Figure SF3**

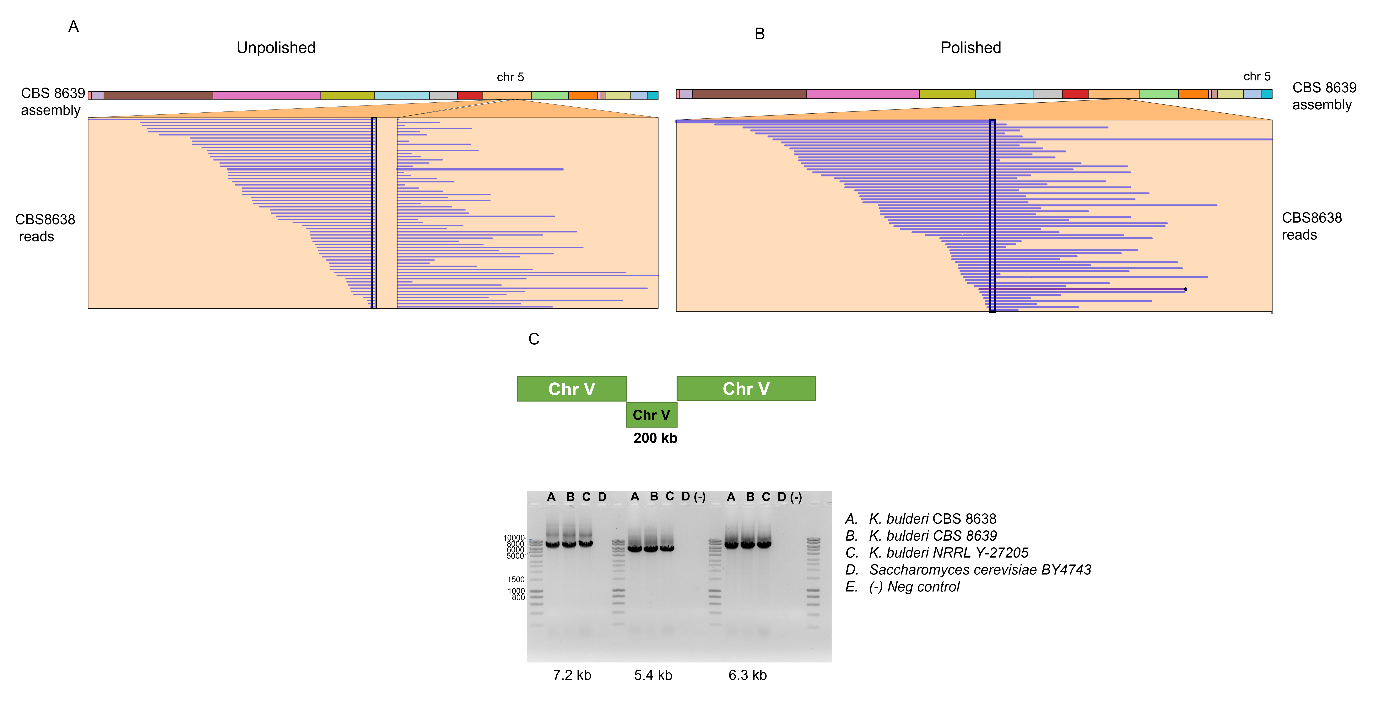

**Figure SF3.** Visual representation of the alignment of CBS 8638 reads against unpolished (Pael A) and polished (Panel B) CBS 8639 genome assembly. Panel C: PCR-based confirmation of the missing region in chromosome V that was identify by using the alternative assembly. Three combinations of primers inside the missing region were used to amplify products in either CBS 8638, CBS 8639 or NRRL Y-27205 strain. *S. cerevisiae* BY4743 was used as control for primer specificity and no-template was used in the control.

**References**

1.	Wick RR, Schultz MB, Zobel J, Holt KE. Bandage: interactive visualization of de novo genome assemblies. Bioinformatics. 2015 Oct 15;31(20):3350–2.

2.	Li H. Minimap2: pairwise alignment for nucleotide sequences. Bioinformatics. 2018;34(18):3094–100.

3.	Li H. New strategies to improve minimap2 alignment accuracy. Bioinformatics. 2021;37(23):4572–4.

4.	Nattestad M, Aboukhalil R, Chin CS, Schatz MC. Ribbon: intuitive visualization for complex genomic variation. Bioinformatics. 2021 Apr 20;37(3):413–5.
