## Supplementary material for "High quality *de novo* genome assembly of non-conventional yeast *Kazachstania bulderi* a new potential low pH production host for biorefineries": Supplementary Tables and Figures.docx

**Table S1.** Growth parameters calculated at different concentration of lactic acid and formic acid. The table shows the mean and standard deviation of the growth rate, doubling time and area under the curve across three biological replicates

| **Strains** | **Conditions** | **Growth rate *r* [h^-1^]** | | | **Doubling Time t_DT_ [h]** | | | **Area under the curve (AUC)** | | |
| --- | --- | --- | --- | --- | --- | --- | --- | --- | --- | --- |
| CBS 8638 | SD+ 50g/L Lactic acid | 0.19 | ± | 0.00 | 3.61 | ± | 0.05 | 108.32 | ± | 3.10 |
| CBS 8639 |  | 0.16 | ± | 0.00 | 4.38 | ± | 0.08 | 99.32 | ± | 5.37 |
| NRRL Y-27205 |  | 0.14 | ± | 0.02 | 5.10 | ± | 0.66 | 110.14 | ± | 5.18 |
| BY4743 |  | 0.03 | ± | 0.00 | 22.13 | ± | 2.73 | 8.54 | ± | 0.72 |
| CBS 8638 | SD+ 60g/L Lactic acid | 0.22 | ± | 0.01 | 3.09 | ± | 0.08 | 113.48 | ± | 2.81 |
| CBS 8639 |  | 0.15 | ± | 0.01 | 4.73 | ± | 0.18 | 104.56 | ± | 5.25 |
| NRRL Y-27205 |  | 0.15 | ± | 0.01 | 4.75 | ± | 0.19 | 98.42 | ± | 10.12 |
| BY4743 * |  | 0.07 | ± | 0.01 | 10.84 | ± | 1.80 | 2.68 | ± | 0.48 |
| CBS 8638 | SD+ 75g/L Lactic acid | 0.32 | ± | 0.01 | 2.14 | ± | 0.09 | 79.13 | ± | 1.47 |
| CBS 8639 |  | 0.30 | ± | 0.02 | 2.35 | ± | 0.17 | 72.99 | ± | 3.54 |
| NRRL Y-27205 |  | 0.20 | ± | 0.00 | 3.43 | ± | 0.05 | 67.18 | ± | 2.10 |
| CBS 8638 | SD+ 80g/L Lactic acid | 0.26 | ± | 0.01 | 2.64 | ± | 0.07 | 77.45 | ± | 2.30 |
| CBS 8639 |  | 0.24 | ± | 0.01 | 2.88 | ± | 0.13 | 71.21 | ± | 2.41 |
| NRRL Y-27205 |  | 0.06 | ± | 0.01 | 12.12 | ± | 2.11 | 25.59 | ± | 8.47 |
| CBS 8638 | SD+ 85g/L Lactic acid | 0.19 | ± | 0.03 | 3.62 | ± | 0.50 | 106.67 | ± | 3.88 |
| CBS 8639 |  | 0.08 | ± | 0.00 | 8.23 | ± | 0.32 | 90.52 | ± | 5.44 |
| NRRL Y-27205 |  | 0.08 | ± | 0.01 | 8.53 | ± | 1.31 | 12.62 | ± | 1.52 |
| CBS 8638 | SD+ 25mM formic acid | 0.50 | ± | 0.01 | 1.40 | ± | 0.02 | 65.50 | ± | 2.43 |
| CBS 8639 |  | 0.47 | ± | 0.01 | 1.48 | ± | 0.04 | 64.22 | ± | 3.29 |
| NRRL Y-27205 |  | 0.43 | ± | 0.02 | 1.60 | ± | 0.06 | 62.30 | ± | 4.04 |

* BY4743 does not grow on lactic acid ≥ 50g/L or/and formic acid ≥ 25mM

**Table S2.** Assembly statistics of raw PacBio *K. bulderi* *de novo* genome assemblies by different approaches IPA and Hifiasm

|  | | **IPA** | | | **Hifiasm** | |
| --- | --- | --- | --- | --- | --- | --- |
|  |  | ***CBS8638*** | ***CBS8639*** | ***NRRL-Y27205*** | ***CBS8639*** | ***CBS8638*** |
| **Primary contig assembly** | Size (Mb) | 14.47 | 14.03 | 14.32 | 17.17 | 19.96 |
|  | N° Contigs | 17 | 14 | 15 | 40 | 26 |
|  | Largest contig (bp) | 2734419 | 2786749 | 2758527 | 3868179 | 3273188 |
|  | N50 (Mb) | 1.37 | 1.12 | 1.10 | 1.51 | 2.44 |
|  | N75 (Mb) | 0.71 | 0.86 | 0.73 | 1.14 | 1.63 |
|  | L50 | 4 | 4 | 4 | 3 | 4 |
|  | L75 | 8 | 7 | 8 | 6 | 6 |
|  | GC (%) | 33.12 | 33.11 | 33.16 | 33.45 | 33.2 |
| **Alternate haplotigs** | Size (Mb) | 13.37 | 14.41 | 16.11 | 10.81 | 7.66 |
|  | N° Contigs | 85 | 108 | 172 | 38 | 33 |
|  | Largest contig (bp) | 1622056 | 2499223 | 1891183 | 1116028 | 1309555 |
|  | N50 (Mb) | 0.92 | 1.13 | 0.95 | 0.67 | 0.69 |
|  | N75 (Mb) | 0.42 | 0.54 | 0.49 | 0.52 | 0.39 |
|  | L50 | 6 | 5 | 7 | 7 | 4 |
|  | L75 | 11 | 9 | 12 | 12 | 8 |
|  | GC (%) | 33.11 | 33.01 | 33.05 | 33.15 | 33.23 |

**Table S3**. Summary statistics of the PacBio reads mapped to the IPA and Hifiams *K.bulderi* assemblies

| **Feature** | **IPA** | | | **Hifiasm** | |
| --- | --- | --- | --- | --- | --- |
|  | **CBS8638** | **CBS8639** | **NRRL Y27205** | **CBS8638** | **CBS8639** |
| Mean read length | 6504.7 | 7083.2 | 7540.8 | 6,191.70 | 6,709.20 |
| Median read length | 6289 | 7042 | 7538 | 6,007.00 | 6,817.00 |
| Number of reads | 131888 | 125890 | 159349 | 145,295.00 | 130,073.00 |
| Read length N50 | 8065 | 8201 | 8927 | 7,950.00 | 8,166.00 |
| Total bases | 857894356 | 891703780 | 1201624508 | 899,629,549.00 | 872,685,868.00 |
| Coverage X | 59.30 | 63.54 | 83.92 | 45.07 | 50.84 |

**Table S4.** |Comparison of the BUSCO statistics on the three *K. bulderi* strains for the Initial and Polished assemblies.

| BUSCO score (%) | | | |
| --- | --- | --- | --- |
|  | ***CBS8638*** | ***CBS8639*** | ***NRRL- Y27205*** |
| Initial assembly | | | |
| Complete | 98.90% | 97.60% | 97.30% |
| Fragmented | 0.30% | 0.30% | 0.30% |
| Missing | 0.80% | 2.10% | 2.40% |
| Polished assembly | | | |
| Complete | 99.00% | 99.10% | 98.70% |
| Fragmented | 0.20% | 0.30% | 0.20% |
| Missing | 0.80% | 0.60% | 1.10% |

**Table S5.** Detailed results of chromosome-level assemblies genome annotation. *Chromosome VIII contains rRNA repetitions

|  | **Chromosome** | **Size (Mb)** | **CDS** | **tRNAs** | **Ty** |
| --- | --- | --- | --- | --- | --- |
| CBS8639 | Chr I | 2.79 | 1093 | 40 | 8 |
|  | Chr II | 2.72 | 1091 | 30 | 4 |
|  | Chr III | 1.42 | 538 | 24 | 3 |
|  | Chr IV | 1.24 | 405 | 19 | 4 |
|  | Chr V | 1.12 | 454 | 13 | 2 |
|  | Chr VI | 0.99 | 380 | 15 | 2 |
|  | Chr VII | 0.86 | 341 | 9 | 0 |
|  | Chr VIII* | 0.80 | 297 | 12 | 4 |
|  | Chr IX | 0.75 | 264 | 14 | 2 |
|  | Chr X | 0.72 | 271 | 18 | 3 |
|  | Chr XI | 0.43 | 145 | 7 | 0 |
|  | Chr XII | 0.40 | 156 | 7 | 3 |
|  | Total | | 5435 | 208 | 35 |
| CBS8638 | Chr I | 2.77 | 1090 | 40 | 7 |
|  | Chr II | 2.73 | 1091 | 30 | 4 |
|  | Chr III | 1.37 | 527 | 24 | 3 |
|  | Chr IV | 1.25 | 488 | 19 | 4 |
|  | Chr V | 1.18 | 460 | 13 | 2 |
|  | Chr VI | 0.94 | 377 | 15 | 3 |
|  | Chr VII | 0.78 | 341 | 9 | 0 |
|  | LB Chr VIII* | 0.80 | 294 | 12 | 5 |
|  | LB Chr IX | 0.73 | 259 | 14 | 2 |
|  | LB Chr X | 0.71 | 266 | 18 | 2 |
|  | Chr XI | 0.43 | 152 | 7 | 1 |
|  | Chr XII | 0.40 | 159 | 7 | 3 |
|  | Total | | 5504 | 208 | 36 |
| NRRL Y-27205 | Chr I | 2.76 | 1083 | 39 | 6 |
|  | Chr II | 2.72 | 1089 | 30 | 6 |
|  | Chr III | 1.43 | 542 | 24 | 3 |
|  | Chr IV | 1.22 | 482 | 19 | 3 |
|  | Chr V | 1.23 | 473 | 13 | 2 |
|  | Chr VI | 1.01 | 380 | 15 | 0 |
|  | Chr VII | 0.90 | 346 | 9 | 0 |
|  | Chr VIII* | 0.84 | 294 | 12 | 11 |
|  | Chr IX | 0.73 | 261 | 14 | 2 |
|  | LB Chr X | 0.65 | 250 | 18 | 3 |
|  | LB Chr XI | 0.55 | 171 | 7 | 0 |
|  | LB Chr XII | 0.40 | 154 | 7 | 1 |
|  | Total | | 5525 | 207 | 37 |

**Table S6.** Number of *K. bulderi* functional annotated proteins using genomes from different yeast species.

| Yeast species | **CBS 8638** | **CBS 8639** | **NRRL Y-27205** |
| --- | --- | --- | --- |
| ***S. cerevisiae*** | 4541 | 4543 | 4523 |
| ***Y. lipolytica*** | 3575 | 3591 | 3578 |
| ***S. pombe*** | 3058 | 3065 | 3062 |
| ***C. albicans*** | 3833 | 3838 | 3819 |
| ***C. glabrata*** | 4395 | 4404 | 4386 |
| ***K. marxianus*** | 4199 | 4211 | 4187 |
| ***K. lactis*** | 4296 | 4298 | 4276 |
| ***K. exigua*** | 5056 | 5069 | 5041 |
| ***K. barnettii*** | 5076 | 5086 | 5049 |

**Table S7**. Genetic variants inside *K. bulderi* strains using CBS 8638 and NRRL Y-27205 as references

| **Reference assembly** | **Reads mapped** | *Heterozygote site* | | | *Outside heterozygote sites* | | | |
| --- | --- | --- | --- | --- | --- | --- | --- | --- |
|  |  | **Same variant combination** | **Different variant** | **Total** | **SNPs** | **Indels** | **Total** | **Total variants** |
| CBS8638 | CBS8639 | SNPs:13 Indels:19 | SNPs:0 Indel:0 | 32 | 716 | 89 | 805 | 837 |
|  | NRRL Y27205 | SNPs:42 Indel:10 | SNPs:0 | 52 | 1247 | 82 | 1329 | 1381 |
|  |  |  | Indel:0 |  |  |  |  |  |
| NRRL Y27205 | CBS8639 | SNPs:5 Indels:9 | SNPs:1 Indel:1 | 16 | 1308 | 109 | 1417 | 1433 |
|  | CBS8638 | SNPs:14 Indel:18 | SNPs:0 | 34 | 1297 | 172 | 1469 | 1503 |
|  |  |  | Indel:2 |  |  |  |  |  |

**Table S8.** Genotype and sources of yeast strains

| **Strain** | | **Genotype and auxotrophic markers** | **Source** |
| --- | --- | --- | --- |
| *S. cerevisiae* | BY4741 | *MATa his3Δ1 leu2Δ0 met15Δ0 ura3Δ0;* [rho ^+^] | Invitrogen |
|  | BY4743 | *MATa/α his3Δ1/his3Δ1 leu2Δ0/leu2Δ0 LYS2/lys2Δ0 met15Δ0/MET15 ura3Δ0/ura3Δ0;* [rho ^+^] | Invitrogen |
|  | KGY029 | *Mata leu2Δ0 leu2-3,112 ura3-52 his3::HindIII arg8::hisG ;* [rho^-^ *COX2*::*ARG8*m] | Cai's lab |
|  | NCYC 505 | prototroph | NCYC^*^  Brewer's top yeast, isolated from beer, Oranjeboom brewery, Rotterdam, Netherlands. |
|  | 96.2 | prototroph | E. Barrio  Natural isolate from *Quercus ilex* bark (Castellón, Spain) |
| *K. bulderi* | CBS 8638 | Prototroph, [rho^0^] (this study) | ^**^ARS culture Collection (NRRL)  Isolated from maize silage |
|  | CBS 8639 | Prototroph, [rho^0^] (this study) | ARS Culture Collection (NRRL)  Isolated from maize silage |
|  | NRRL Y-27205 | Prototroph, [rho^-^ ] (this study) | ARS Culture Collection (NRRL)  Isolated from maize silage |

^*^NCYC: National Collection of Yeast Cultures

^**^Agricultural Research Service culture collection (Northern Regional Research Laboratory)

**Table S9.** Primer sequences for verification of inversion

| **Name** | **Sequence** | **Amplicon Size** |
| --- | --- | --- |
| F1 | CCGTTGACCTATGACTGTAC | 1787 |
| R1 | GATCGCCTGTGGTAAATCTG |  |
| F2 | CACCAAGAGCCTGGATTAGA | 3041 |
| R2 | GCGAATTTAACTTCACCTGG |  |
| F1 + F2 |  | 1596 |
| R1 + R2 |  | 2363 |

**Supplementary Figures**

**Figure S1**

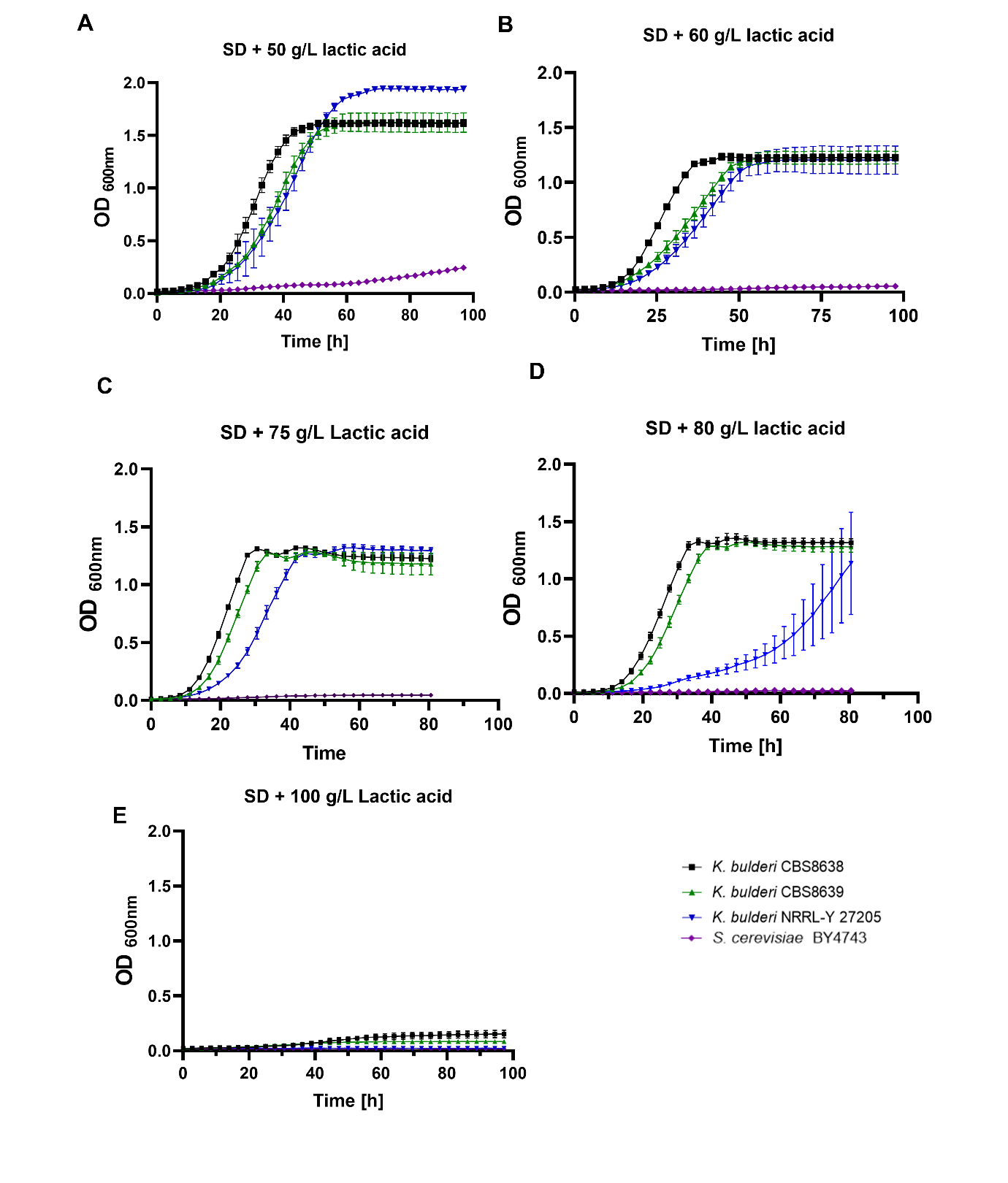

**Figure S1**. Growth curves of *K. bulderi* CBS 8638, CBS 8639 and NRRL Y-27205 strains in A. 50 g/L B. 60 g/L and C. 75 g/L D. 80g/L and E. 100 g/L of lactic acid at 2.5 pH.

**Figure S2**

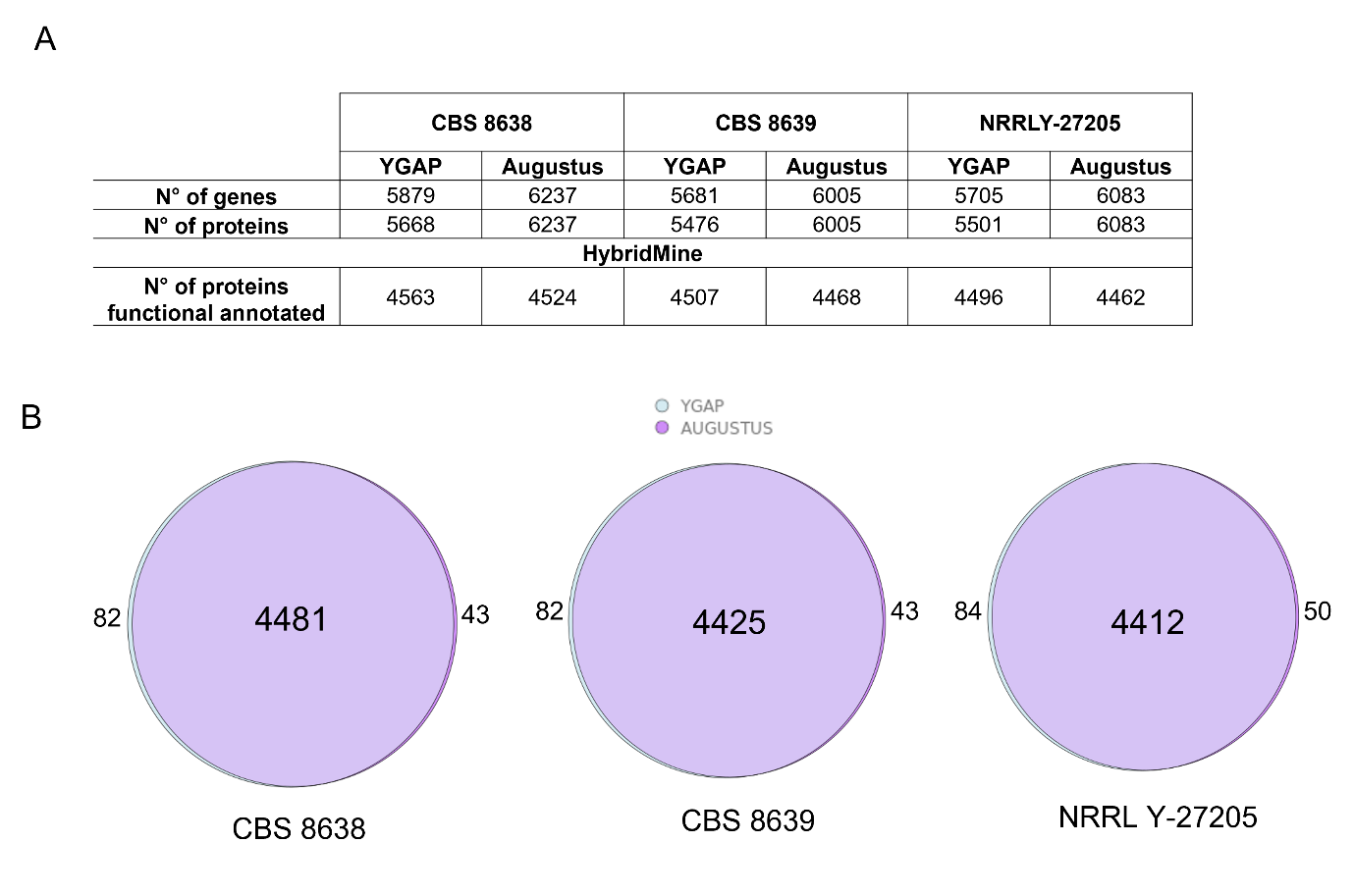

**Figure S2**. A. Table containing the number of genes and proteins annotated by YGAP and AUGUSTUS for IPA assemblies of CBS 8638 and CBS 8639, and NRRL Y-27205, and the number of proteins predicted for HybridMine when using *S. cerevisiae* as reference. B. Proportional Venn diagram with the number of functional annotated proteins predicted for HybridMine in common between the methods for structural annotation, YGAP light blue and AUGUSTUS dark purple, YGAP&AUGUSTUS light purple.

**Figure S3**

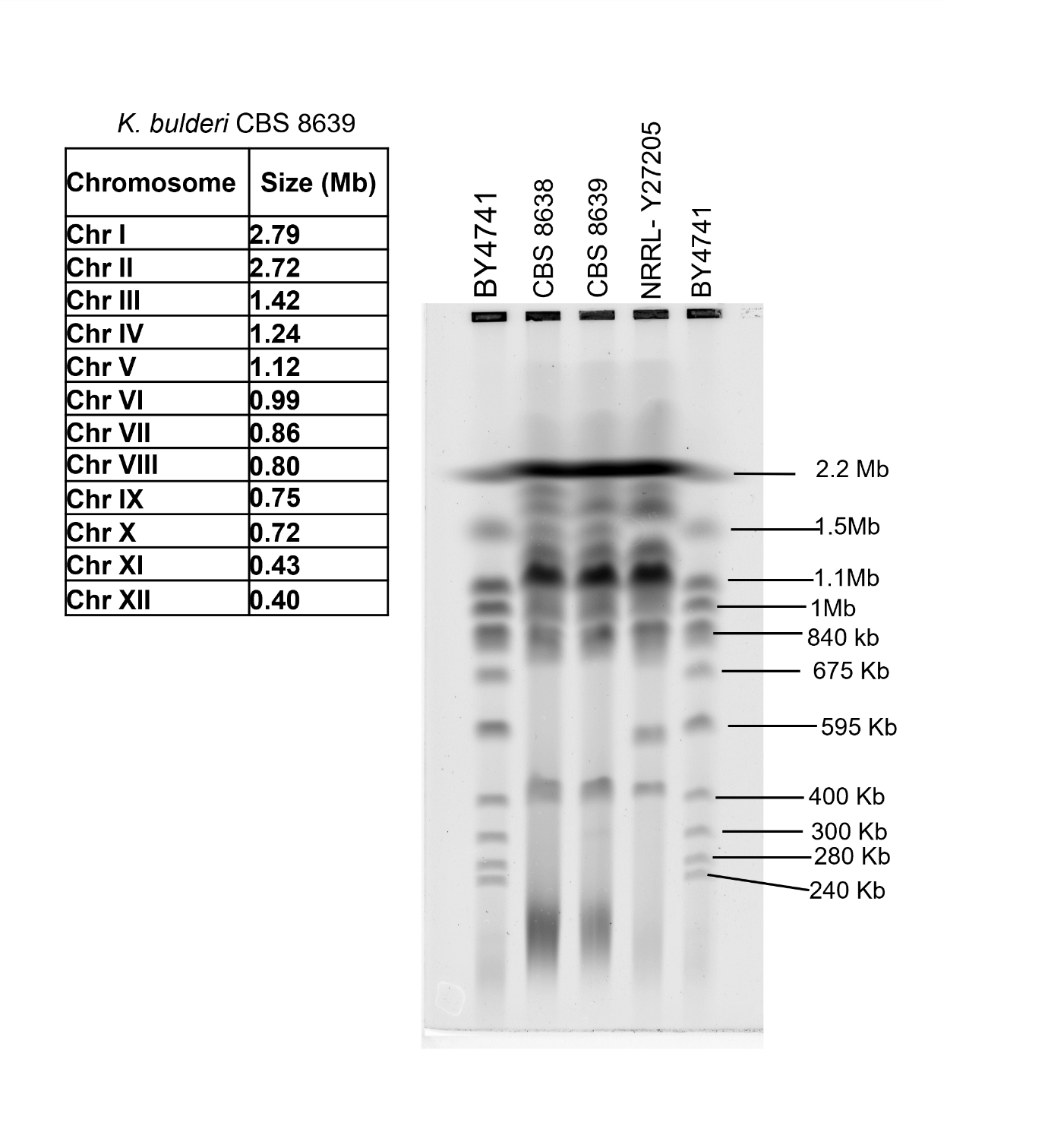

**Figure S3.** Analysis by Pulse Field Gel Electrophoresis (PFGE) of chromosomal DNA from the *Kazachstania* *bulderi* CBS 8638, CBS 8639 and NRRL Y-27205 strains. The first and the last line contain S. cerevisiae BY4741 strain as size marker.

**Figure S4.**

**A.**
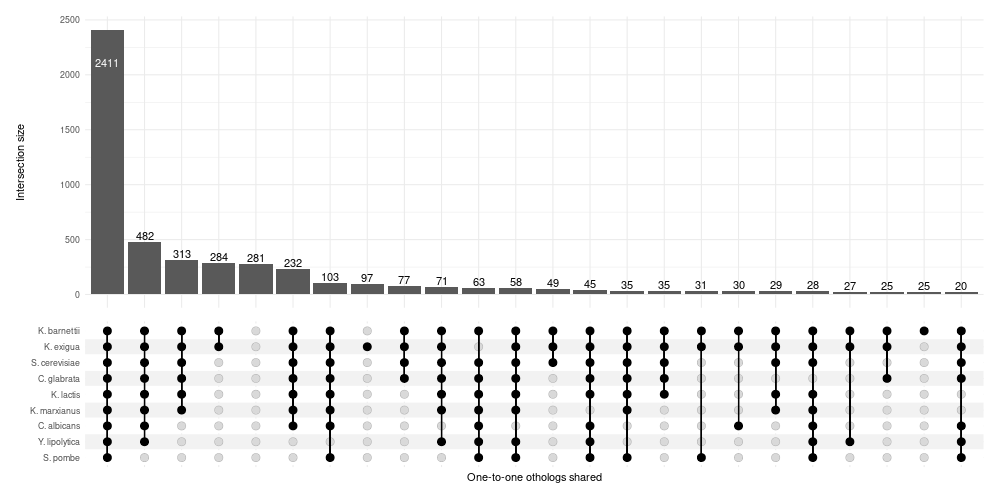

**B.**

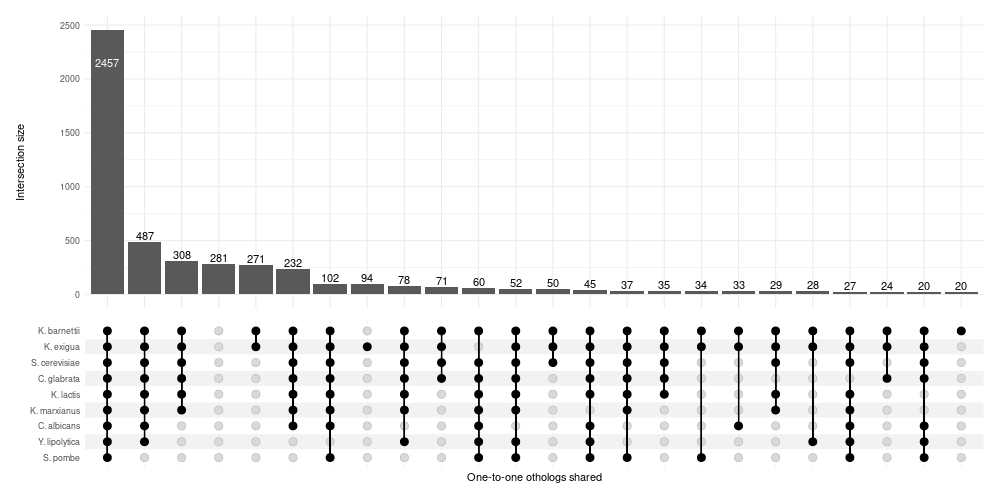

**Figure S4.** Upset plot showing the number of proteins functionally annotated across *K. bulderi* A. CBS 8638 and B. NRRL Y-27205 strains in common when using *Saccharomyces cerevisiae*, *Schizosaccharomyces pombe, Candida albicans,* *Candida glabrata*, *Yarrowia lipolytica*, *Kluyveromyces marxianus,* *Kluyveromyces lactis, Kazachstania exigua* and *Kazachstania barnettii*, as references genomes. Vertical lines across the species represent the proteins annotated in common among them. Individual dots represent the proteins of each species shared only with *K. bulderi*.

Figure S5

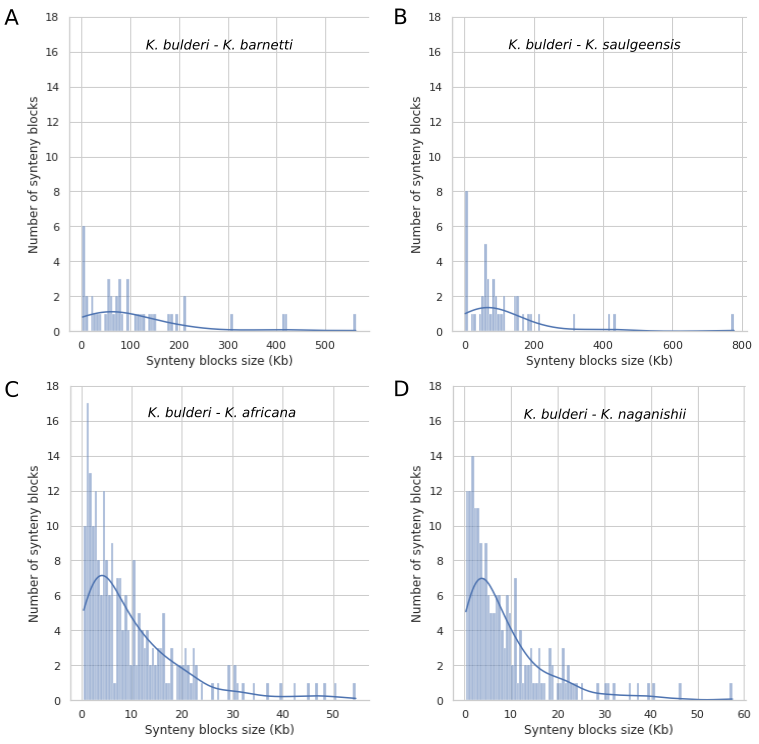

Figure S5. Histogram representing the occurrence of the synteny blocks sizes between *K. bulderi* CBS 8639 strain and *K. barnettii* (Panel A), *K. saugeensis* (Panel B), *K. africana* (Panel C) and *K. naganishii* (Panel D).

Figure S6

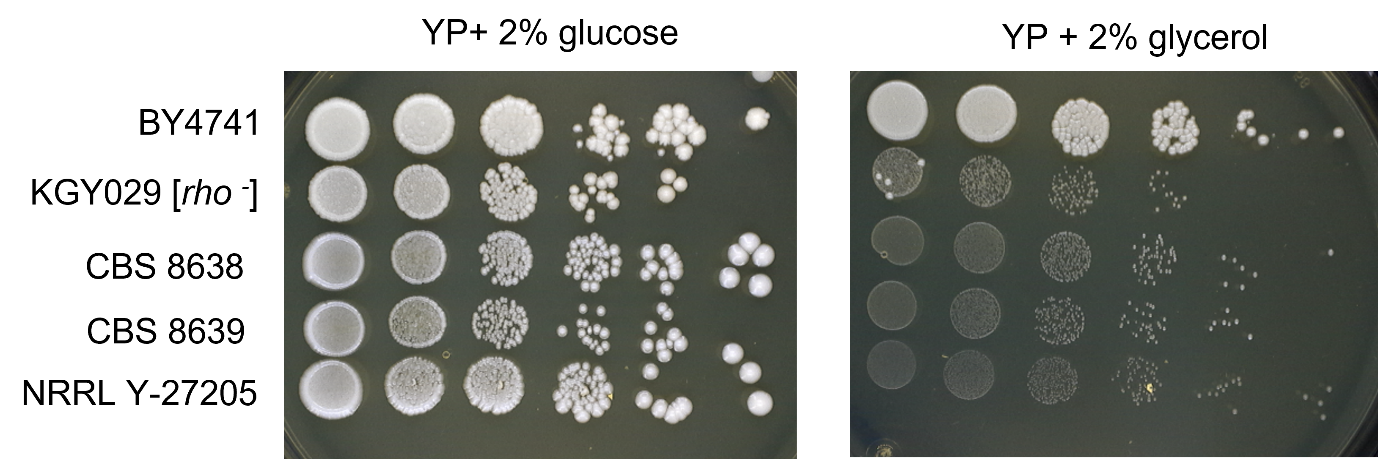

Figure S6. Spot test assay for CBS 868, CBS 8639 and NRRL Y-27205 *K. bulderi* strains on YP+ 2% glucose and YP+ 2% glycerol. Comparison is made against *Saccharomyces cerevisiae* BY4741 (WT) and COX2-mutant rho ^–^ strain contains a mutation on *COX2* gene affecting mtDNA and avoid its grow in glycerol.
